# Molecular characterization of thermo-adapted Novirhabdovirus variants and virulence *in vivo* in rainbow trout and zebrafish at different temperatures

**DOI:** 10.64898/2026.10.29.762795

**Authors:** Calvin Fauvet, Elise Gourhannic, Manon Mehraz, Julie Bernard, Annie Lamoureux, Mélissa Aurousseau, Emilie Mérour, Maxence Frétaud, Lénaïg Louboutin, Yannick Blanchard, Thierry Morin, Dimitri Rigaudeau, Christelle Langevin, Jean K. Millet, Stéphane Biacchesi

## Abstract

Controlling viral diseases in fish is a challenge for the sustainable development of aquaculture and improving the health of fish farms. Infectious hematopoietic necrosis virus (IHNV) is a major pathogen in aquaculture. It infects cold-water fish, but loses its infectivity at temperatures above 20°C. Since fish are poikilothermic animals, their body temperature fluctuates according to that of their environment. Faced with current climate change, it is crucial to understand how these pathogens will adapt to rising temperatures. Through a strategy simulating the impact of global warming, this virus has been progressively thermo-adapted to 25°C. The selected variant, IHNV_25_, replicates at 25°C while maintaining efficient replication at low temperatures, like its parental strain. Genome sequencing and a reverse genetics approach revealed that only 9 amino acid substitutions, distributed across 4 of the 6 viral genes, were essential for thermo-adaptation. Although IHNV_25_ was highly attenuated in its natural host, the rainbow trout, at any temperatures, it proved highly virulent when injected in zebrafish larvae at 20°C and 25°C, inducing severe symptoms identical to those observed in trout (hemorrhages, oedema, and exophthalmia). At both temperatures, mortality was preceded by a massive viral replication and a delayed and inefficient interferon and inflammatory response. Unlike the parental strain, IHNV_25_ was also capable of infecting larvae by bath immersion at 25°C, entering via the caudal fin, thus demonstrating its ability to cross the species barrier. The identification of viral markers of thermo-adaptation will therefore be of great interest for the epidemiological surveillance of field isolates.

## Introduction

Climate change poses an unprecedented existential threat to organized human life, with grave consequences for virtually all living species, ecosystems, and the biosphere. This is a key and timely issue, given that 2023 broke almost every climate record and saw what was likely the highest average of daily global temperature recorded on Earth in 100,000 years [1]. The steady rise in sea and ocean surface temperatures is particularly concerning (Copernicus Climate Change Service), with global and North Atlantic surface temperatures reaching new highs in 2023. Thus, one of the main challenges of the coming decades will consist of feeding more than 9 billion people by 2050, in the context of climate change.

Global fish consumption continues to rise, and future demand cannot be met by capture fisheries alone. Aquaculture is the fastest-growing food sector in the world (6% annually) and currently constitutes the primary source of aquatic food products [2]. Approximately 94% of freshwater fisheries take place in developing countries, providing food and livelihoods for millions of the world’s poorest people. The development of sustainable aquaculture must overcome several challenges: ensuring animal health and welfare, combating emerging infectious diseases, and adapting farming practices to societal demands and environmental crises, such as human-induced changes, pollution, and global environmental shifts. Consequently, controlling pathogens that affect farmed fish, without compromising environmental commitments, is a major priority. This requires evolving farming practices, limiting antibiotic use, and developing effective vaccines suitable for mass administration [3].

Warming is the primary abiotic factor influencing the physiology of ectothermic organisms, such as fish. Temperature alters molecular processes, cellular structures, and the rates of biochemical reactions; it thereby affects fish metabolism, growth, and key physiological functions, such as sex determination, sexual maturation, stress responses, and immune system function [4]. As poikilothermic animals, fish have an internal temperature that varies with that of their environment. Chronic exposure to high temperatures imposes a physiological and metabolic burden that can impair the ability of fish to cope with other stressors and, consequently, compromise their survival [5]. These temperature variations can be substantial and serve as a major factor modulating immune activity [6]. While low temperatures limit immune competence, temperatures exceeding the tolerance range can lead to immunosuppression. Thus, temperature fluctuations can have either negative or positive effects on fish resilience and their ability to mount an appropriate response to infections, depending on how these variations impact their immune competence.

Climate model projections indicate that global surface temperatures, across both land and surface waters (oceans and rivers), could rise by 0.4 to 2.6°C by 2050 [7, 8]. Optimal and lethal temperatures (both minimum and maximum) vary significantly among fish species. Rainbow trout (*Oncorhynchus mykiss*) is one of the most highly valued salmonid species worldwide, owing to its commercial value, rapid growth rate, and history of introduction across multiple continents. Depending on the strain, rainbow trout can survive in water temperatures ranging from 0°C to 27°C; however, the preferred temperature range for optimal physiological performance is between 15°C and 17°C [9]. In certain geographic regions, water temperatures may approach the limits of the optimal range for some currently farmed species. Consequently, climate change represents a major challenge that will significantly impact aquaculture production and its sustainability, due to fish migration and the probable emergence of fish viruses and new variants, combined with warming waters, hypoxia, increased CO_2_ concentrations, and acidification.

It is important to consider that both fish hosts and pathogens are affected by variations in water temperature. Among fish pathogens notifiable by the WOAH, infectious hematopoietic necrosis virus (IHNV; species *Novirhabdovirus salmonid*) causes acute diseases in salmonids (trout and salmon) in major production region: Europe, Asia, and North America. This EU regulated pathogen is recognized as the cause of one of the most serious diseases affecting finfish in the Northern Hemisphere. Recent epidemiological data show an increase in IHNV outbreaks in rainbow trout farms, accompanied by an expansion of its geographical distribution in Europe [10, 11]. IHNV is a rhabdovirus belonging to the *Novirhabdovirus* genus. With an optimal *in vivo* replication temperature between 4 to 15°C, the virus primarily affects juveniles but can also infect fish of all sizes and ages, resulting in mortality rates ranging from 5 to 90% [12]. At higher water temperatures (15 to 19°C), IHNV-induced disease is associated with lower mortality rates, and no mortality is observed above 20°C. Like most RNA viruses, it possesses a relatively compact genome (11 kb) encoding five structural proteins (the nucleoprotein N, the phosphoprotein P, the matrix protein M, the glycoprotein G and the RNA-dependent RNA polymerase L) as well as one non-structural protein (non-virion protein NV) [13]. The viral polymerase is intrinsically error-prone with estimated substitutions rates for *Novirhabdovirus* species ranging from 2×10^−3^ to 4×10^−5^ substitutions/site/year [14]. In a preliminary study of IHNV outbreaks in an Idaho region characterized by intensive aquaculture, Troyer and Kurath suggested that higher water temperatures (approximately 15°C) were associated with a three- to four-fold increase in viral evolutionary rates [15]. A similar pattern was suggested for another cold-water *Novirhabdovirus* species, the viral hemorrhagic septicemia virus (VHSV; species *Novirhabdovirus piscine*), where higher temperatures in freshwater farms compared to marine environments was associated with higher replication rates and a 2.5-fold increase in the nucleotide substitution rate in the viral glycoprotein (G) gene [16]. These characteristics enable *Novirhabdovirus* species to rapidly adapt to new environmental conditions and hosts. Consequently, they serve as excellent viral models for studying the effects of rising temperatures on their evolution, adaptation, virulence, and host range.

In fact, the emergence of naturally thermo-adapted viral strains has already been reported and could pose a serious threat in the event of epidemics [17, 18]. Furthermore, although most *Novirhabdovirus* strains circulating in salmonids are unable to replicate *in vitro* or *in vivo* at temperatures above 20°C, thermo-adapted VHSV and IHNV variants were successfully obtained through *in vitro* selection involving repeated cell culture passages ([19] and Monique LeBerre, unpublished data). It is worth noting that the *Novirhabdovirus* genus already includes species adapted to temperatures exceeding 30°C, such as the snakehead rhabdovirus (*Novirhabdovirus snakehead*), isolated in Southeast Asia. Documented increases in the replication of viruses affecting fish and mammals have been linked to rising temperatures. This phenomenon is likely associated with higher mutation rates, potentially leading to accelerated evolution and altered fitness profiles [20, 21]. In fish, increased virulence associated with rising temperatures has already reported for viral nervous necrosis in Japan [22]. More recently, Combe and colleagues predicted that a 1°C rise in water temperature would lead to an increase in mortality rates, ranging from 1% to 8%, for major viral pathogens in aquaculture, including Cyprinid Herpesvirus 3 and viral nervous necrosis virus [23].

The hypothesis that global warming exerts significant selective pressure on aquatic viruses and acts as a major stressor for host fish raises the question of the emergence of thermo-adapted viral strains exhibiting modified virus-host interactions, particularly regarding host range and pathogenicity. It is essential to consider these temperature-induced changes in virus-host interactions to improve surveillance and strengthen aquaculture resilience. This study aimed to examine the effects of global warming on IHNV, a major viral pathogen of salmonids with high evolutionary potential. Specifically, the main objectives were to address the following questions: 1) What are the molecular determinants of thermo-adapted IHNV strains? 2) What are the effects of rising temperatures on *Novirhabdovirus* evolution of host range?

## Results

### Selection and proteome analysis of thermo-adapted IHNV variants using a global warming simulator

In the 1980s, a Canadian strain of IHNV (IHNV-CAN67, genotype M [24]), hereinafter referred to as IHNV_WT_, was provided to our laboratory by D. F. Amend (University of California Davis School of Veterinary Medicine, Davis, CA). Originally isolated in 1967 from a diseased rainbow trout (*Oncorhynchus mykiss*) in British Columbia, the strain was subsequently adapted through 78 successive *in vitro* passages in a cyprinid-derived cell line (EPC) at progressively increasing temperatures. This process, carried out using a global warming simulator shown in Figure 1, resulted in the selection of a thermo-adapted variant, IHNV_25_ (Monique LeBerre, unpublished data). This variant has acquired the capacity to replicate efficiently *in vitro* at 25°C, which is 5°C above the replication temperature limit of its parental IHNV_WT_ strain. Briefly, the thermal adaptation process was performed on EPC cells to ensure the viability of the cell at 25°C, given that salmonid-derived cell lines have an optimal growth limit at 20°C. At that time, a large number of passages, combined with a gradual increase in temperature, were performed with the aim of empirically selecting an attenuated variant suitable for use as a live attenuated vaccine. Thus, the parental IHNV_WT_ underwent 7 passages at 19°C, 25 at 21°C, 26 at 23°C and 20 at 25°C (Monique LeBerre, unpublished data).

**Fig. 1.**
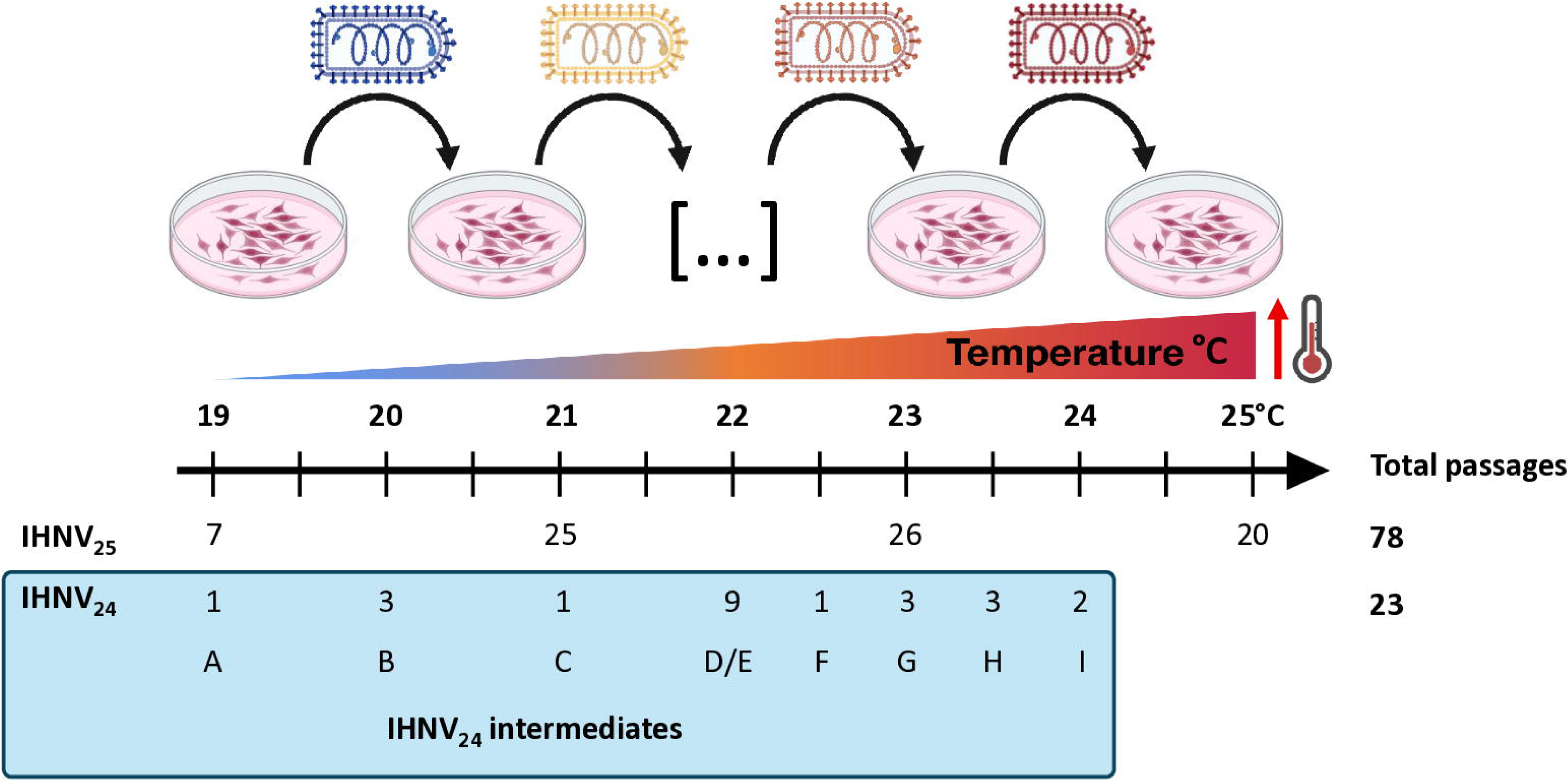
*In vitro* viral evolution in global warming simulator. Schematic representation of the thermo-adaptation of two parental IHNV strains to increasing temperatures in EPC cell culture, indicating the number of passages at each temperature stage. For variant IHNV_24_-mCherry (subsequently referred to as IHNV_24_), the stabilized intermediates (#A to I, excluding #D) obtained at each temperature are indicated and were fully sequenced. At 22°C, two intermediates were analyzed: #D, obtained after a single passage, and the stabilized intermediate #E, obtained after 8 additional passages. The images were adapted from BioRender.

Since IHNV_25_ variant was recovered after 78 successive passages in EPC cells, a second thermo-adapted variant has recently been obtained using a similar *in vitro* protocol simulating global warming. A recombinant IHNV-mCherry (rIHNV-mCherry [25]), expressing a red-fluorescent protein and derived from the French IHNV strain 32/87 (isolated from rainbow trout in France in 1987 and also belonging to genotype M), was adapted to 24 °C through only 23 passages in EPC cells. This limited number of passages was performed in order to restrict the number of mutations related to adaptation to cell culture (Fig. 1). Thus, the parental IHNV 32/87 strain underwent 1 passage at 19°C, 3 at 20°C, 1 at 21°C, 9 at 22°C, 1 at 22.5°C, 3 at 23°C, 3 at 23.5°C and 2 at 24°C. The process was stopped at 24°C following the loss of the red-fluorescent protein. Importantly for this variant, subsequently referred to as IHNV_24_, all intermediates were conserved. This will make it possible to track the chronological order in which thermo-adaptive substitutions appeared in the genome.

The genomes of parental strains and thermo-adapted variants were then fully sequenced and compared (Fig. 2 and Tables 1 and 2; GenBank: IHNV CAN67, QB032804; IHNV 32/87, QB032805; IHNV_25_, QB032806 and IHNV_24_-mCherry, QB032807). Analysis of the IHNV_25_ genome revealed that a limited number of eleven amino acid substitutions had appeared during the thermal adaptation process and were sufficient to confer it the ability to replicate efficiently at 25 °C (Fig. 2A and Table 1). These substitutions were distributed across the various viral proteins, with the exception of the NV protein, for which no amino acid changes were observed. In addition, no substitutions were detected in non-coding regions (3’ leader, 5’ trailer and intergenic regions). Similarly, analysis of the IHNV_24_ genome revealed that only nine amino acid substitutions had appeared in viral proteins during this more rapid adaptation process (Fig. 2B and Table 2). These substitutions were distributed across the various viral proteins, with the exception of the P and the NV proteins, for which no amino acid changes were observed. An additional nucleotide substitution was detected in the 5’ trailer region (Table 2). Four mutations were also observed in the mCherry gene that could explain the fluorescence extinction of the mCherry protein at 24°C: three resulting in drastic amino acid changes and another causing the loss of the stop codon and the extension the ORF beyond the gene end signal of the expression cassette (Fig. 2). The acquisition of these substitutions took place chronologically during the different stages of temperature increase, as illustrated in Figure 2B. Once selected, they were retained throughout the adaptation process to 24°C. It is interesting to note that, for the two thermo-adapted variants, some of these mutations were identified in the NCBI GenBank database via BLAST searches on various field isolates, primarily within the glycoprotein (Tables S1 and S2).

**Fig. 2.**
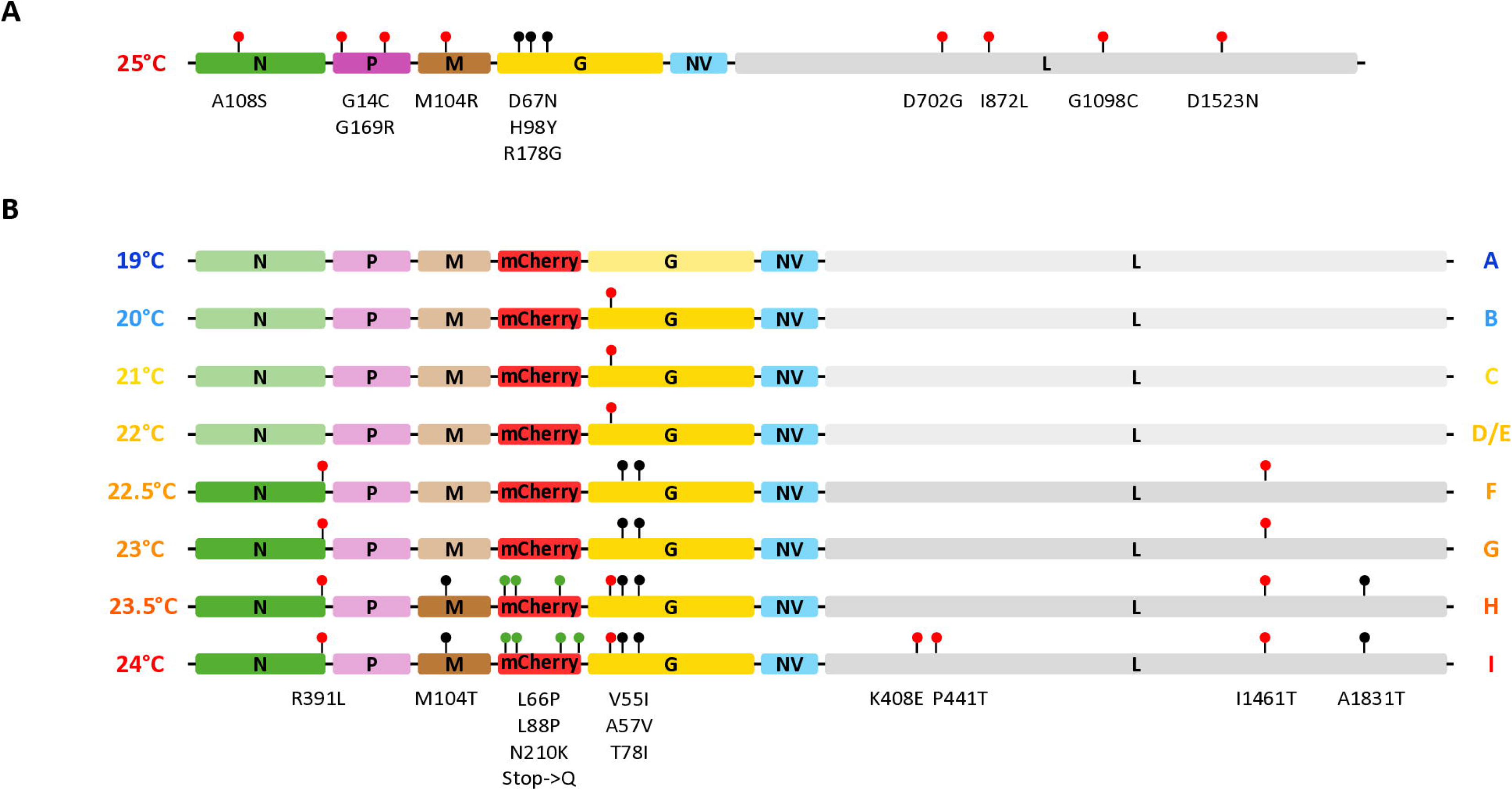
Genetic mapping of substitutions identified in the thermo-adapted strains IHNV_25_ (A) and IHNV_24_ (B). For each virus, the sequence of each viral protein was compared by amino acid alignment between the parental strain and the corresponding thermo-adapted variant. Markers indicate the mutations identified between the thermo-adapted variants and the parental strains. Black markers correspond to amino acid polymorphisms observed in other field strains, whereas red markers correspond to novel variations. For IHNV_24_, the sequences of the intermediates (#A to I) at the indicated temperature are presented, illustrating the appearance of each mutation over successive passages at increasing temperatures. Green markers correspond to mutations accumulated in the mCherry protein.

**Table 1:**
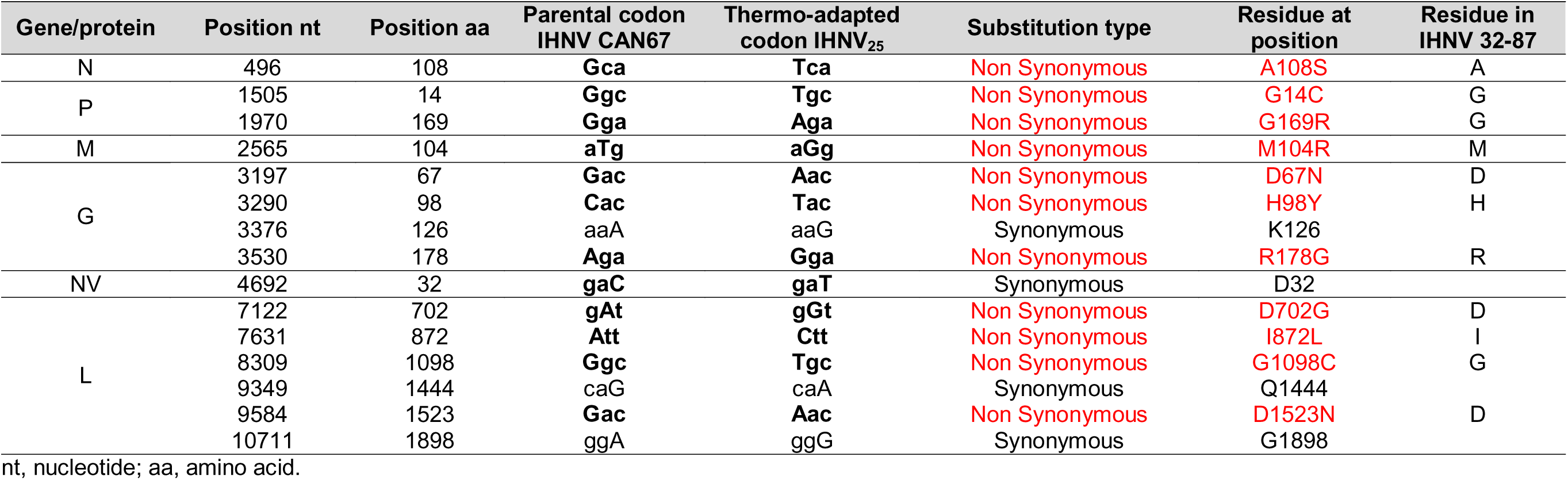
Mutations identified in the thermo-adapted IHNV_25_ compared to its parental IHNV CAN67.

**Table 2:** Mutations identified in the thermo-adapted IHNV_24_ compared to its parental IHNV 32-87.

| Gene/protein | Position nt<sup>a</sup> | Position aa | Parental codon<br>IHN<sub>V</sub>32-87 | Thermo-adapted<br>codon IHN<sub>V</sub><sub>24</sub> | Substitution type | Residue at<br>position | Residue in<br>IHN<sub>V</sub> CAN67 |
| --- | --- | --- | --- | --- | --- | --- | --- |
| N | 651 | 159 | agC | agT | Synonymous | S159 |  |
|  | 1346 | 391 | cGc | cTc | Non Synonymous | R391L | R |
| M | 2573 | 104 | aTg | aGg | Non Synonymous | M104T | M |
| G | 3169 | 55 | Gtc | Atc | Non Synonymous | V55I | V |
|  | 3176 | 57 | gCc | gTc | Non Synonymous | A57V | A |
|  | 3239 | 78 | aCa | aTa | Non Synonymous | T78I | T |
|  | 3474 | 156 | aaC | aaT | Synonymous | N156 |  |
| L | 6246 | 408 | Aaa | Gaa | Non Synonymous | K408E | K |
|  | 6345 | 441 | Ccc | Acc | Non Synonymous | P441T | P |
|  | 9406 | 1461 | aTa | aCa | Non Synonymous | I1461T | I |
|  | 10515 | 1831 | Gcg | Acg | Non Synonymous | A1831T | A |
| Trailer | 11058 | — | T | C | — | — |  |

<sup>a</sup> nucleotide position in the genome of IHN<sub>V</sub> 32-87 strain. nt, nucleotide; aa, amino acid.

**Table 3:**
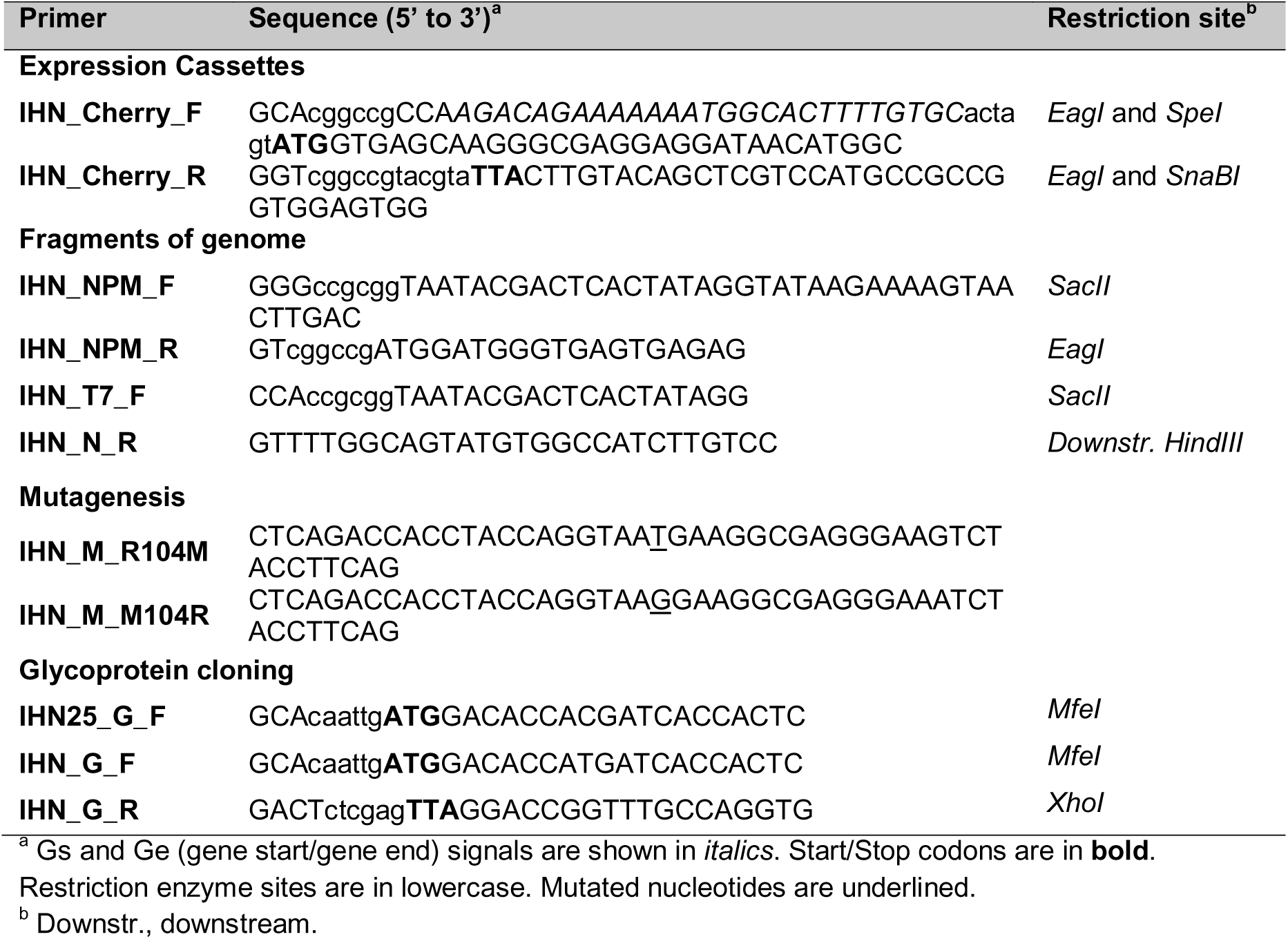
Primers for cDNA constructs.

**Table 4.**
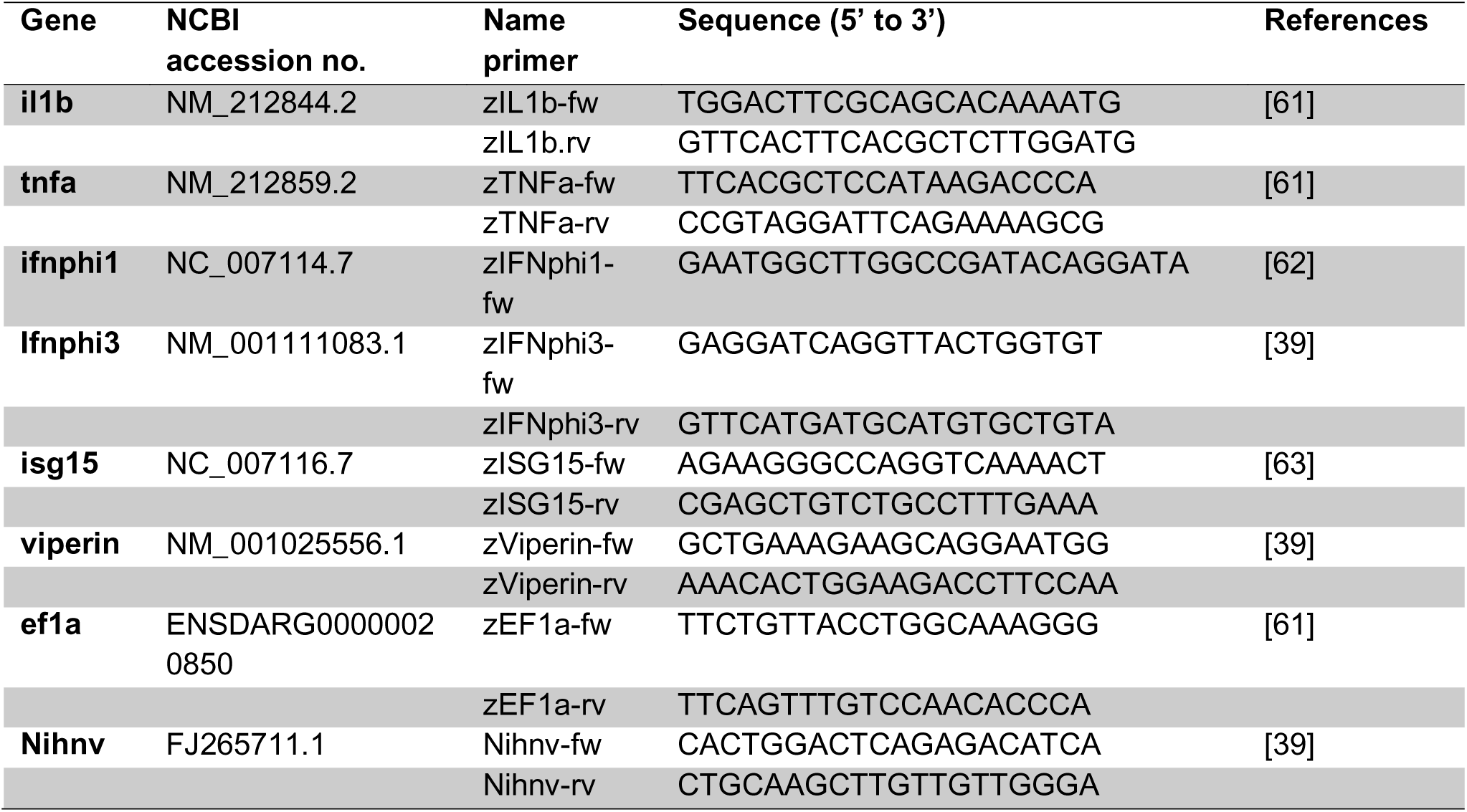
Primers to evaluate zebrafish gene expression and viral replication.

The substitutions found in thermo-adapted strains were mapped onto AlphaFold-predicted three-dimensional structures of the N, P, M, G, and L IHNV parental strain proteins (Fig. 3). Each of the predicted IHNV proteins adopt a structure similar to known rhabdovirus protein structures [26]; [27]; [28]; [29]. N is predicted with good overall confidence (pTM 0.62-0.64) and organized in two lobes composed mainly of alpha helices (N-terminal and C-terminal domains) with an extended C-terminal tail lacking secondary structure, likely indicative of a disordered region similar to other rhabdovirus N proteins (Fig. 3A) [26]. For IHNV_25_, a substitution (A108S) was found in the N-terminal domain, in a region distal from the putative RNA binding groove of the protein (Fig. 3A, left). For IHNV_24_, the last residue of the protein was found to be substituted (R391L), at the tip of the non-structured C-terminal tail of the protein (Fig. 3A, right). Both mutations are surface exposed and could be involved in modulating interactions for N oligomerization, or with other components of the replication machinery (e.g. P or L proteins), or with host cell factors.

**Fig. 3.**
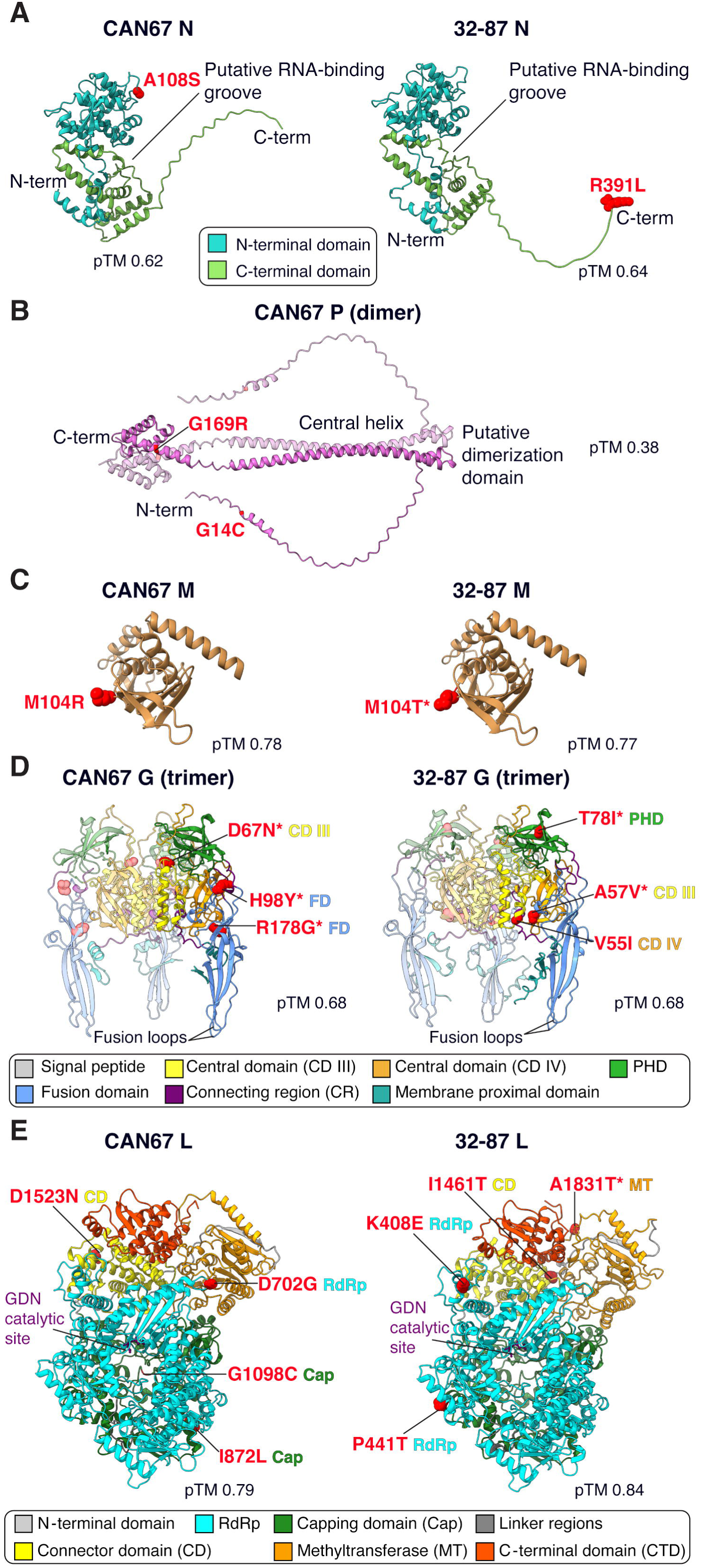
Mapping of substituted sites found in thermo-adapted strains on the predicted structures of IHNV CAN67 and 32-87 parental strains. IHNV_25_ and IHNV_24_ thermo-adaptation substitutions were mapped onto AlphaFold 3-predicted protein structures of the respective parental strains, IHNV CAN67 and 32-87. (A) Nucleoprotein, N; (B) Phosphoprotein, P modeled as dimer; (C) Matrix protein, M (structured globular domain shown); (D) Glycoprotein, G, modeled as trimer (ectodomains shown); and (E) Large protein (viral RNA-dependent RNA polymerase), L. The sites where substitutions occur are highlighted in red, with parental residues shown in atom sphere representation. Asterisks denote substitutions that have been previously identified in field strains. The predicted template modelling (pTM) scores are shown for each predicted structure. Protein structure labeling and representation were visualized using ChimeraX.

Based on the known quaternary structure of rabies virus (RABV) P that forms dimers, the IHNV P was predicted as a dimer (Fig. 3B) [26]. AlphaFold 3 predicts that the protein mostly folds in alpha helical secondary structures with a long, mostly disordered N-terminal region, an extended central helix forming an axis for the P dimer, and a C-terminal domain organized in a bundle of alpha helices. The disordered N-terminal region likely explains the relatively low prediction score for CAN67 P (pTM 0.38). The two substitutions found for IHNV_25_ both involve glycine residues, with G14C situated at the beginning of a small alpha helix within the mostly disordered N-terminal region and G169R located on an alpha helix in the C-terminal region. For RABV, the disordered P N-terminal domain is involved in mediating binding to N0, the monomeric nucleoprotein unbound to RNA while the P C-terminal domain is involved in binding to RNA-bound multimeric N proteins. Thus, the substitutions found in IHNV_25_ may affect N-P interactions.

The predicted M protein for both parental strains fold with high confidence (pTM 0.78 and 0.77) into a globular domain containing both alpha helices and beta strands similar to other rhabdoviruses such as the vesicular stomatitis virus (VSV) (Fig. 3C) [27]. Interestingly, the substitution found in IHNV_25_ (M104R) and IHNV_24_ (M104T) involve the same residue position (M104) in their respective parental strains, CAN67 and 32-87. The site appears surface exposed and may be implicated in interactions such as M-M multimerization or interactions with other viral (N) or host cell proteins. The substitution M104T was identified in other field isolates (Table S2).

As rhabdovirus G proteins form homotrimers, the IHNV predicted proteins were modeled as trimers [30]. AlphaFold generated trimeric models of IHNV G with good confidence scores (pTM 0.68) that adopt a pre-fusion conformation for both parental strains (Fig. 3D). For each thermo-adapted strain, a set of 3 distinct substitutions were identified distributed across several domains. For IHNV_25_, a substitution was found in the central domain III region (CD III, D67N) near the G protein central helix. This substitution involves a change from an aspartic acid (D) to an asparagine (N) residue, the former having a negatively charged side chain while the latter having a polar uncharged side chain. The other two substitutions, H98Y and R178G are located in the fusion domain in a region distally located from the membrane-interacting fusion loops.For IHNV_24_, the substitutions involve different G domains with V55I and A57V located in central domains III and IV, respectively. The third substitution, T78I, is located in the pleckstrin homology domain (PHD), near the apex of the trimer in the pre-fusion conformation and the site of major neutralizing determinants and conformational epitopes. With the exception of IHNV_24_ V55I, all G proteins substitutions were identified previously in other field strains (Table S1 and S2). T78I is especially noteworthy since it has been previously been characterized in an IHNV neutralization escape variant with altered tissue tropism and virulence [31]. Overall while the set of mutations found in IHNV_25_ and IHNV_24_ are distinct, they could potentially impact receptor binding or conformational changes required for the fusion function of G, neutralization by antibodies, and/or protein stability and expression at higher temperatures.

The viral polymerases L of CAN67 and 32-87 were modeled with very high confidence pTM scores of 0.79 and 0.84, respectively (Fig. 3E). The overall predicted structures for IHNV L proteins are remarkably similar to the Cryo-EM determined structure of VSV [29]. A set of 4 distinct substitutions were identified in the L protein of each thermo-adapted strains and distributed across several functional domains. For IHNV_25_, the D702G substitution was found in the RdRp domain, located in a loop adjacent to the methyltransferase domain and not in close proximity to the GDN catalytic site. Two substitutions, I872L and G1098C were located in the capping domain, but not in close proximity of each other. A fourth substitution, D1523N, was found in the connector domain. For IHNV_24_, a distinct set of 4 substitutions were mapped involving two residue positions in the RdRp, K408E and P441T, one position in the connector domain (I1461T), and a residue position located in the methyltransferase domain (A1831T), the latter being found in other field strains (Table S2). No common mutations were found between the two thermo-adapted strains. The substitutions identified in the L protein could be implicated in maintaining stability of the protein and/or stabilize interactions with other viral co-factors such as N and P proteins.

### Comparison of replication kinetics and plaque sizes of parental strain and thermo-adapted variant in cell culture at different temperature

A reverse genetics system was established for the IHNV_25_ variant, as previously described [32]. The recombinant IHNV_25_ (rIHNV_25_) replicated as efficiently as the *in vitro* thermo-adapted IHNV_25_ variant at 25°C. The characteristics of thermo-adapted IHNV in cell culture were then studied by comparison with those of IHNV_WT_. Specifically, the morphologies of the plaques produced by rIHNV_25_, and IHNV_WT_ at 15°C and 25°C were compared using a plaque assay (Fig. 4A). The plaques induced by rIHNV_25_ at 15°C were slightly smaller than those of IHNV_WT_, although the difference was not statistically significant (Fig. 4B). In contrast to IHNV_WT_, which does not replicate at 25°C, rIHNV_25_ replicated efficiently at 25°C producing significantly larger plaques than those obtained at 15°C, suggesting enhanced intercellular spread or increased cytopathic activity.

**Fig. 4.**
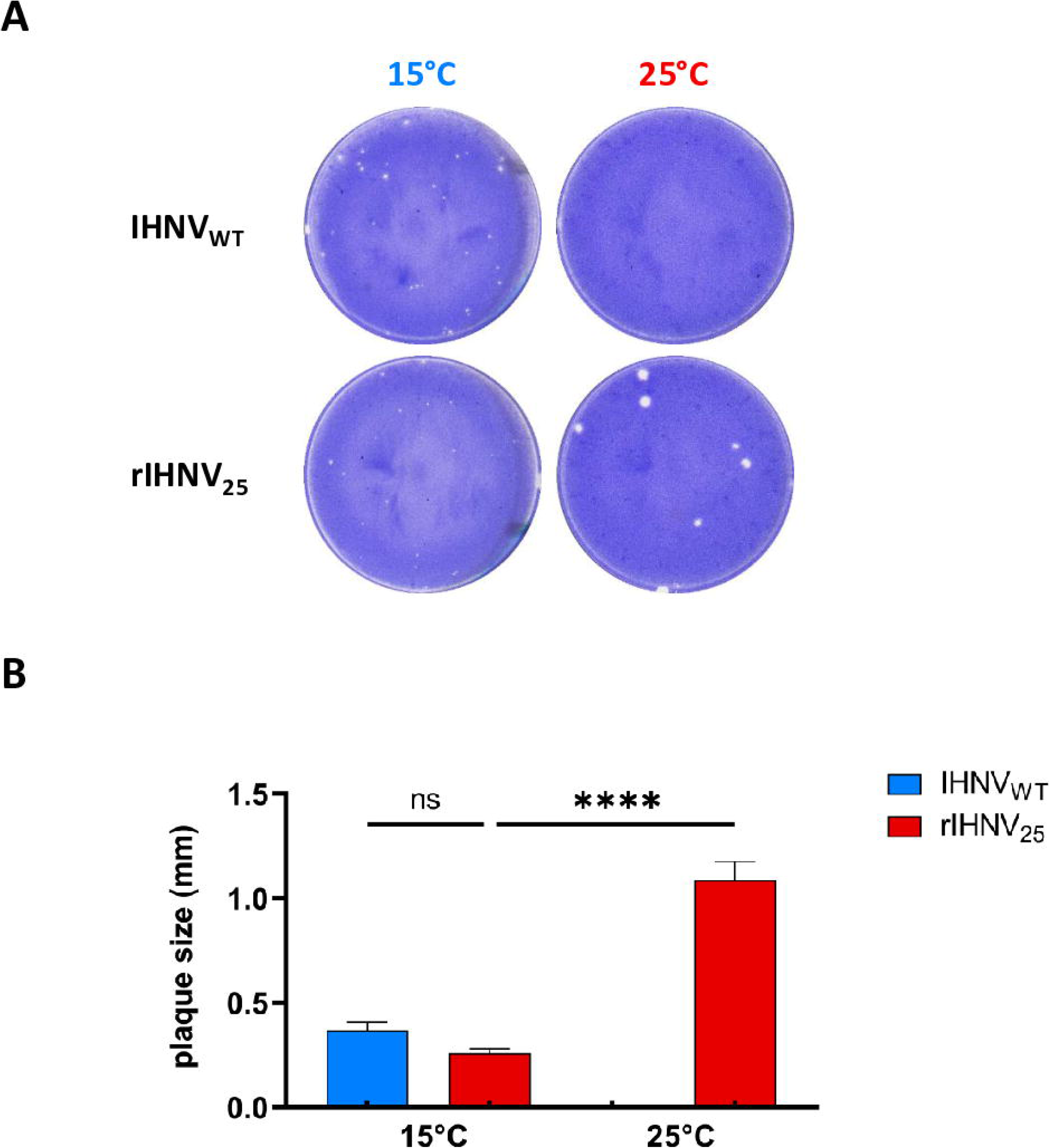
Comparison of growth and plaque size between the parental virus and the thermo-adapted recombinant virus rIHNV_25_ generated by reverse genetics at two temperatures. **A.** Growth ability and plaque morphology of the parental virus (wild-type IHNV; IHNV_WT_) and the thermo-adapted recombinant virus (rIHNV_25_), generated by reverse genetics, were evaluated at 15°C and 25°C. EPC cells were infected with 20 to 40 PFU of each virus and cultured in medium containing Avicel. At 3 days post infection, cell monolayers were fixed and stained. **B.** Comparison of plaque sizes between the parental virus and the thermo-adapted recombinant virus at 15°C and 25°C. Means are presented with standard errors. Groups showing no significant difference are marked ns (not significant), and those showing a significant difference are marked **** (p < 0.0001).

Comparison of the multi-step growth curves of IHNV_WT_, IHNV_25_ and rIHNV_25_, following infection of EPC cells at various temperatures with a constant multiplicity of infection of 0.01 (Fig. 5A), showed that the viral production and replication kinetics of IHNV_25_ and rIHNV_25_ were strictly identical at all four temperatures tested, thereby validating the IHNV_25_ genome sequence and the reverse genetics system. Although adapted to growth at higher temperatures, rIHNV_25_ was nonetheless capable of replicating efficiently at low temperatures, specifically 10°C and 15°C, with only a very slight delay at 10°C compared to IHNV_WT_. In contrast, at 20°C, the replication kinetics of rIHNV_25_ was much faster than that of IHNV_WT_. rIHNV_25_ reached a plateau at day 3 post infection, with a viral yield of roughly 5.5×10^7^ PFU/mL, a value 10-fold higher than that reached by IHNV_WT_. As expected, IHNV_WT_ did not replicate at 25°C, whereas rIHNV_25_ reached a peak of replication at day 3 post infection, with a final titer of 8×10^6^ PFU/mL, slightly lower than that obtained at 20°C.

**Fig. 5.**
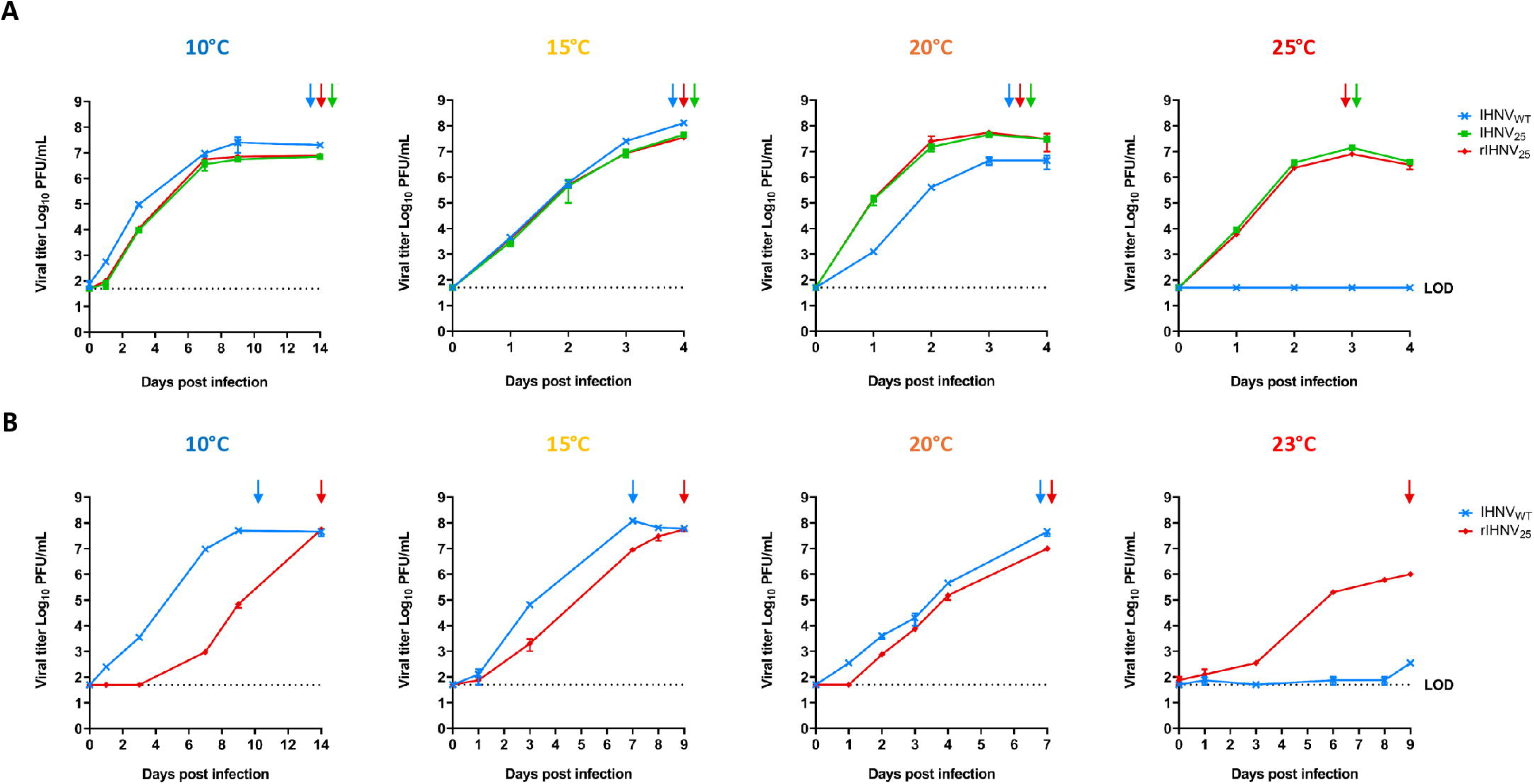
Multi-step growth curves at different temperatures in EPC and RTgill-W1 cells. EPC (A) and RTgill-W1 (B) cells were infected at an MOI of 0.01 PFU per cell with the parental strain (IHNV_WT_), the thermo-adapted variant (IHNV_25_) and the thermo-adapted recombinant IHNV (rIHNV_25_) and then incubated at 10°C, 15°C, 20°C, 23°C and 25°C. Supernatant aliquots were harvested on the indicated days post infection and subsequently analyzed by plaque assay. Each virus titration was performed in duplicate. Mean values are presented with the calculated standard errors. Arrows indicate the time point at which a total cytopathic effect was observed. The limit of detection (LOD) for virus titration is 5 × 10^1^ PFU/mL.

Similar multi-step growth curves were established for IHNV_WT_ and rIHNV_25_ following infection of RTgill-W1 cells at various temperatures, using a constant multiplicity of infection of 0.01 (Fig. 5B). RTgill-W1 cell line was selected because it is derived from rainbow trout gill and gill cells have previously been shown to serve as a suitable cellular substrate for distinguishing the virulence phenotypes of field isolates [33]. Indeed, in this cell line, rIHNV_25_ exhibited a much more pronounced phenotype compared to that observed in EPC cells and that of IHNV_WT_. Although it reached a final titer similar to that of IHNV_WT_, rIHNV_25_ production was delayed by 1 to 3 days depending on the temperature. The difference in replication kinetics was considerably reduced when the cells were incubated at 20°C. At 23°C, the upper temperature limit for this cell line, the rIHNV_25_ variant proved fully capable of replicating in trout-derived cells, unlike its parental strain. The rIHNV_25_ variant reached a final titer of 1 × 10 PFU/mL on day 9 post-infection, a value 10-fold lower than that obtained at 20°C.

### Characterization of viral genes involved in the thermo-adaptation

Functional studies on recombinant viruses were conducted to identify viral genes involved in adaptation to 25°C, using chimeric constructs based on the exchange of genes or group of genes between the thermo-sensitive strain IHNV 32/87 and the thermo-adapted variant rIHNV_25_ (derived from IHNV CAN67), using the reverse genetics systems available for both viruses. The IHNV CAN67 and IHNV 32/87 strains are closely related and belong to genotype M (> 99% protein pair identity; see Table S3). More importantly, the amino acid at the position of the substitutions identified in rIHNV_25_ are conserved between these two strains (Table 1). The strategy, shown in Figure 6, initially involved the exchange in the rIHNV_WT_ 32/87 backbone of genes previously suspected of being linked to temperature sensitivity, especially the glycoprotein G and polymerase L genes, either individually or in combination [34, 35]. These three recombinant viruses, rIHNV_WT_-G_25_, rIHNV_WT_-L_25_ and rIHNV_WT_-GL_25_, were successfully recovered, with a final titer comparable to that of the parental viruses at 15°C; however, these chimeric viruses failed to replicate at 25°C, similarly to rIHNV_WT_. The nucleoprotein N, the phosphoprotein P and the matrix protein M genes were then successively added to the rIHNV_WT_-GL_25_ backbone, enabling the ready recovery of rIHNV_WT_-NGL_25_, rIHNV_WT_-NPGL_25_ and rIHNV_WT_-NPMGL_25_ viruses, respectively. These three chimeric viruses replicated efficiently at 15°C, however only rIHNV_WT_-NPMGL_25_ was able to replicate at 25°C. In order to validate the necessity of all 5 genes, the glycoprotein G, the polymerase L, the nucleoprotein N and the phosphoprotein P genes were successively replaced by those of the thermo-sensitive strain and rIHNV_WT_-NPML_25_, rIHNV_WT_-NPMG_25_, rIHNV_WT_-MGL_25_ and rIHNV_WT_-NMGL_25_, respectively, were produced and amplified at 15°C. rIHNV_WT_-NMGL_25_ was the only one capable of replicating at 25°C, demonstrating that the minimal combination of genes involved in thermo-adaptation of IHNV included the genes encoding the nucleoprotein N, the matrix protein M, the glycoprotein G and the polymerase L. However, the size of the plaques produced by rIHNV_WT_-NMGL_25_ was slightly smaller than that of the plaques produced by rIHNV_25_ and rIHNV_WT_-NPMGL_25_, indicating that the amino acid substitutions in the phosphoprotein P confer a certain replication advantage at 25°C.

**Fig. 6.**
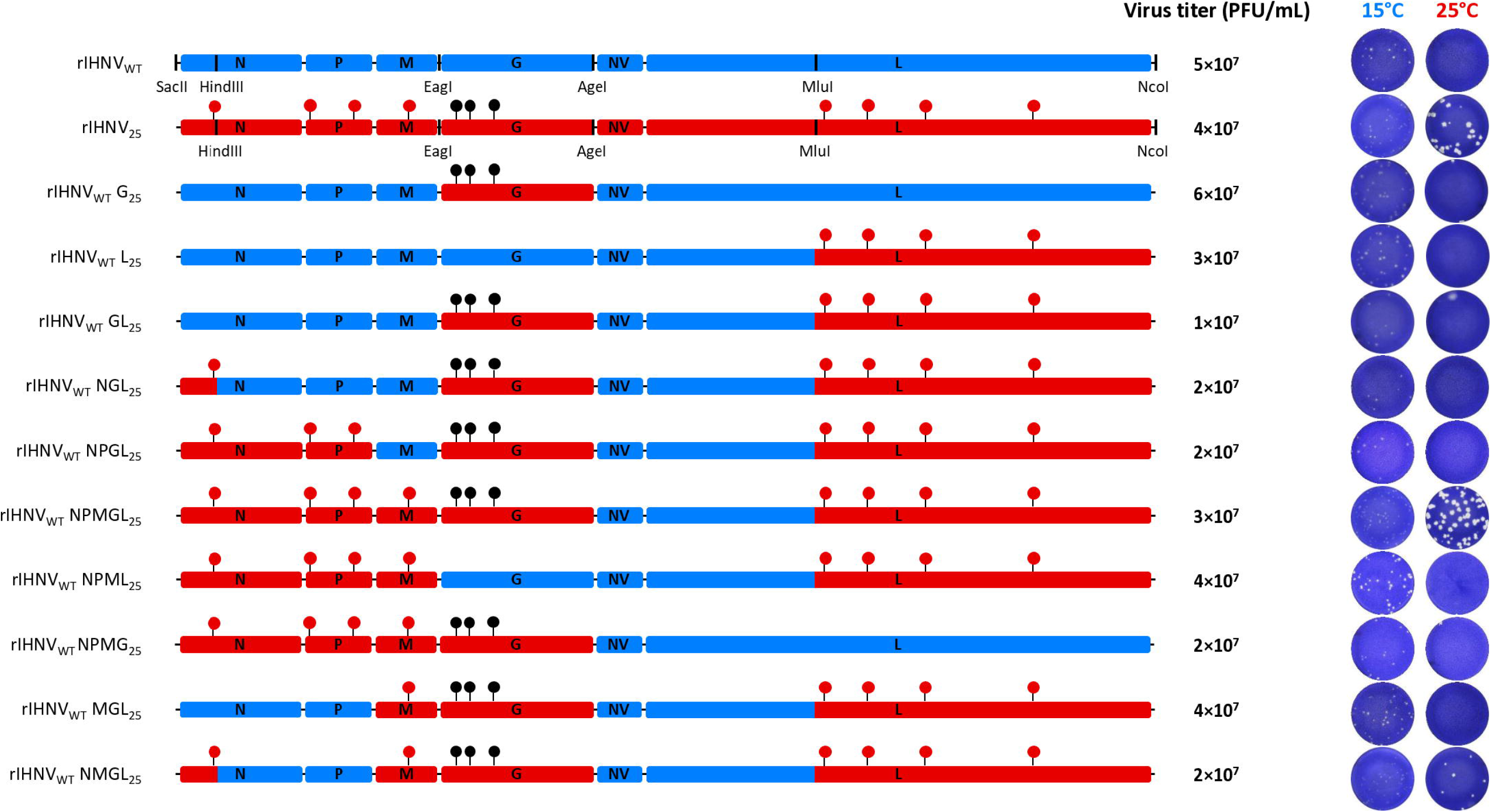
Construction, recovery, and plaque morphology of chimeric recombinant viruses obtained by exchanging genome fragments between the parental strain and the thermo-adapted IHNV variant. The complete parental genome of rIHNV is shown in blue at the top drawn approximately to scale, followed by that of rIHNV_25_, in red, on which are indicated the amino acid substitutions existing between both viruses. The restriction enzyme sites used for the constructs are indicated. Below are the chimeric genome constructs created to define the minimal set of genes required for thermo-adaptation. On the right, the viral titers reached by the various viruses at 15°C are listed, along with the sizes of the plaques observed of the chimeric viruses in EPC cells at 15°C and 25°C, 4 days post infection.

### Expression and targeting at the cell membrane of the glycoprotein G derived from the parental strain and the thermo-adapted variant at 15°C and 25°C

Based on our approach using chimeric viruses, substitutions in the G glycoprotein proved essential, but not sufficient for thermal adaptation. Accordingly, G protein expression and plasma membrane targeting were evaluated in EPC cells transfected with plasmid vectors encoding the G protein derived from either the IHNV_WT_ strain or the IHNV_25_ variant, and maintained at 15°C or 25°C. The expression of G was assessed by indirect immunofluorescence on fixed cells, using a mAb recognizing the unfolded form of G, or on live cells, using a neutralizing mAb recognizing a conformational epitope (Fig. 7A). In transfected and fixed EPC cells, both G proteins were expressed in the cytoplasm, regardless of the incubation temperature. However, at the none-permissive temperature of 25°C, the staining pattern for IHNV_WT_ G protein differed markedly, as it seemed to be retained in the endoplasmic reticulum (ER). In contrast, at 15°C, the IHNV_WT_ G seemed to be distributed throughout the Golgi complex and the plasma membrane, as was observed for IHNV_25_ G protein at both temperatures.

**Fig. 7.**
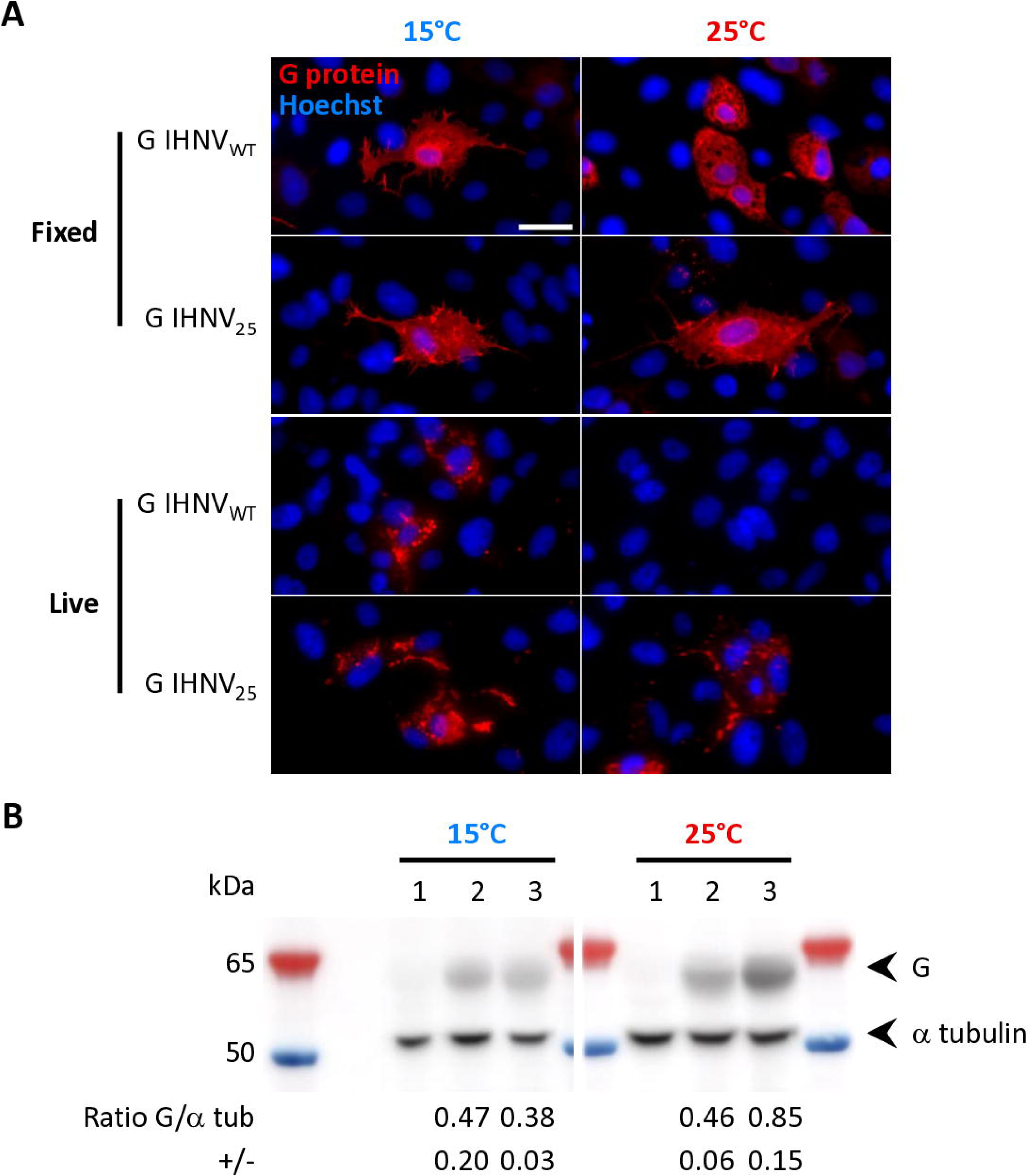
Expression and targeting of the glycoprotein G at the cell membrane from parental and thermo-adapted strains at 15°C and 25°C. **A.** The expression of IHNV glycoprotein derived from the parental (G IHNV_WT_) and the thermo-adapted (G IHNV_25_) strains was evaluated using indirect immunofluorescence assays on EPC cells incubated at 15°C and 25°C. Cells were transfected with plasmid vectors encoding these glycoproteins and incubated at 15°C or 25°C. At 48 hours post transfection, cells were fixed and permeabilized and G protein expression was detected using a mAb against IHNV G protein (red). Membrane expression of G protein was visualized on live cells using another mAb against IHNV G protein (red). Nuclei were stained with Hoechst (blue). Scale bars, 20 µm. **B.** Lysates from transfected cells, harvested at 48 hours post transfection, were analyzed by Western-blot: empty vector (lane 1), IHNV_WT_ G (lane 2) and IHNV_25_ G (lane 3). Proteins were separated by SDS-PAGE and then transferred on a PVDF membrane. IHNV G protein expression was detected using a mAb against IHNV G protein, and alpha-tubulin expression levels were determined as loading controls. Molecular weight markers are shown, with molecular masses in kilodaltons (kDa) indicated on the left. Below each lane is the expression ratio of IHNV G protein and alpha-tubulin, as calculated by the Western-blot imaging system (means of two independent experiments).

We analyzed then the expression of the G proteins on live EPC cells transfected with the same plasmid constructs to verify the correct targeting of the protein to the plasma membrane (Fig. 7A). In accordance with results obtained on fixed samples, at the none-permissive temperature of 25°C, the IHNV_WT_ G was not detected to the surface of EPC cells, unlike the IHNV_25_ G protein. It cannot be excluded that the IHNV_WT_ G at 25°C was misfolded and, consequently, was no longer recognized by the neutralizing mAb.

To further assess whether variations in expression levels contributed to the observed differences, we analyzed the abundance of both G proteins in transfected EPC cells at the two temperatures by Western blot. As shown in Figure 7B, the expression of each G protein was somewhat similar at 15°C. In contrast, at 25°C, the amount of IHNV_WT_ G protein was considerably reduced compared to that of IHNV_25_ G protein, suggesting the degradation of a portion of the misfolded IHNV_WT_ G protein and/or a higher stability of the IHNV_25_ G protein.

### Pathogenicity of the thermo-adapted variant IHNV_25_ in juvenile trout reared at three different temperatures

The *in vivo* pathogenicity of the IHNV_25_ variant in rainbow trout has never been tested, regardless of the temperature. Thus, the pathogenicity of IHNV_WT_, IHNV_25_ and rIHNV_25_ was first evaluated by infecting juvenile rainbow trout (INRAE AUT*SYN reference line) by bath immersion and rearing at 10°C, a temperature at which acute epidemics occur under natural conditions [36]. The virus-induced mortality rate was daily monitored during 28 days post infection (Fig. 8A). For IHNV_WT_ infected group, mortalities began at day 6 post infection and reached a cumulative percent of mortality (CPM) of 94% at day 20, highlighting the high susceptibility of juvenile trout to this IHNV strain under these experimental conditions. Interestingly, no mortality was recorded for IHNV_25_ and rIHNV_25_ throughout the experimental period, indicating that these thermo-adapted variants were completely attenuated in rainbow trout. This attenuation is consistent with the delayed replication of rIHNV_25_ observed in RTgill-W1 cells at 10°C (Fig. 5B), and in contrast with the phenotype observed in EPC cells (Fig. 5A). Fish initially infected with the rIHNV_25_ strain were subsequently challenged with the virulent IHNV_WT_ strain on day 28 post-primary infection (Fig. 8B). Although the 28 days older trout were less susceptible, due to their increased body weight, mortality was observed in the control group (fish that received a mock inoculation) challenged with IHNV_WT_ with a CPM of 35% at day 29 post challenge. Surprisingly, in the group primarily infected with the rIHNV_25_ variant, no significant difference (p > 0.05) was observed in fish mortality kinetics compared to the mock-infected fish group, meaning that the initial infection with the rIHNV_25_ variant did not induce any immune protection.

**Fig. 8.**
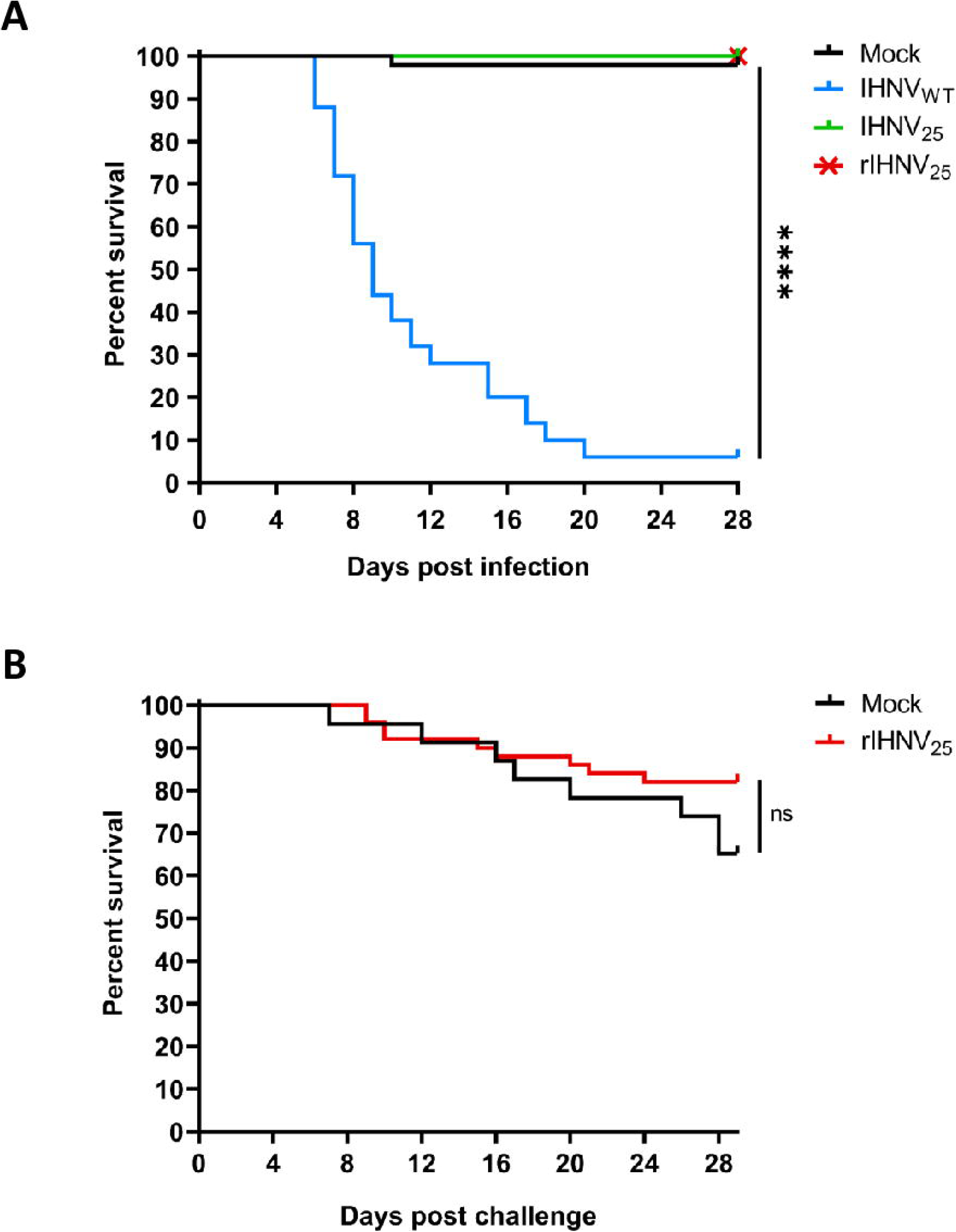
***In vivo* phenotypes of the thermo-adapted IHNV_25_ in rainbow trout reared at 10 °C. A.** Percentage of survival of trout infected by the parental (IHNV_WT_), the thermo-adapted (IHNV_25_) and the thermo-adapted recombinant (rIHNV_25_) viruses, and reared at 10°C. Juvenile rainbow trout (50 fish/tank; mean weight of 0.54 g) were infected by bath immersion with 5 × 10^4^ PFU/mL of each virus or mock infected with cell culture medium as a negative control (Mock). Mortalities were recorded every day for 28 days. **B.** At day 28 post infection, surviving trout from groups infected with rIHNV_25_ or mock-infected were challenged with 5 × 10^4^ PFU/mL of the parental IHNV_WT_ (IHNV CAN67) by bath immersion. Mortality was recorded every day for the following 29 days. Groups that are not significantly different from each other are noted ns (p > 0.05), whereas those that are significantly different are noted ****(p < 0.0001).

To further evaluate the pathogenicity of the thermo-adapted variant at temperatures above 10°C, juvenile rainbow trout (“maigre” line) reared at 10°C were divided in 3 groups. One group of trout was maintained at 10°C as a reference temperature for IHNV infection, while the other two were gradually acclimated to 15°C and 23°C, as shown in Figure 9A. The highest temperature tested of 23°C was chosen to limit the impact on the trout welfare, as trout stop feeding at a temperature of 25°C. Given that higher body weight was associated with better survival following IHNV infection [37], the fish were weighted at the start of the experiment, at the end of the acclimation period, and at the end of the infection trial (only the fish from the mock groups at each temperature). At the start of the experiment, the average weight of the fish was 0.64 g. At the end of the acclimation period, the fish reared at 10°C and 15°C had similar average weights of 0.91 g and 0.94 g, respectively, whereas the trout acclimated at 23°C showed a lower average weight of 0.76 g, reflecting the reduced appetite and the associated growth performance of the trout linked to the rise in temperature. Overall, the acclimatation process was well tolerated by the trout, and no mortality was observed.

**Fig. 9.**
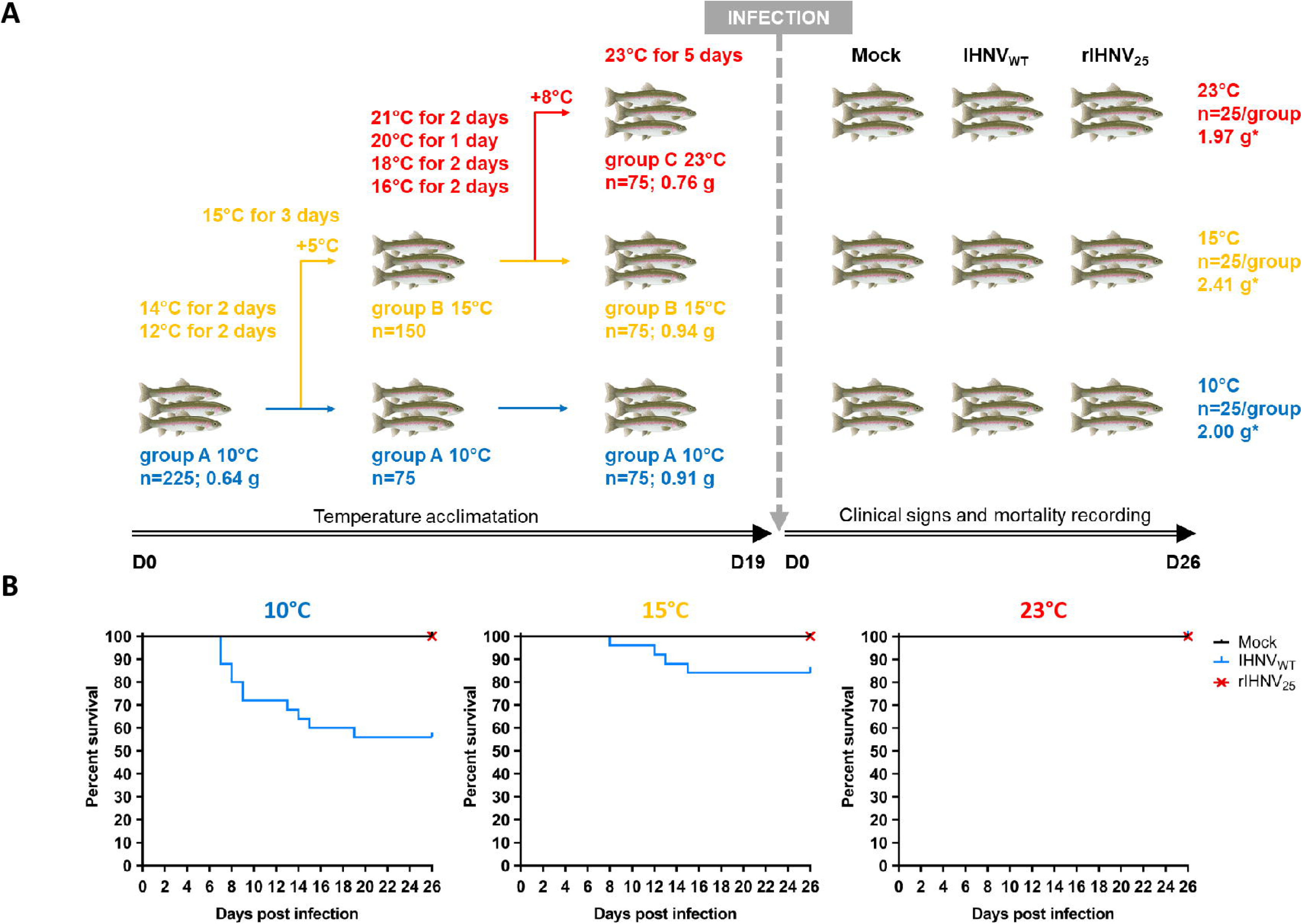
***In vivo* phenotypes of the thermo-adapted IHNV in rainbow trout reared at three different temperatures. A.** Acclimatation protocol for trout at 15°C and 23°C. 225 juvenile trout reared at 10°C were divided in three groups. One group of 75 trout was maintained at 10°C, while the other two groups of 75 trout each were gradually acclimated to 15°C and 23°C, as indicated. These three groups were then subdivided into subgroups of 25 fish and infected by bath immersion with 5 × 10^4^ PFU/mL of the parental strain IHNV_WT_ and the thermo-adapted recombinant variant rIHNV_25_ or mock infected with cell culture medium (Mock) as a negative control. Fish were weighted at the start and the end of the acclimatation protocol as well as at the end of the infection (* only for fish in the Mock groups). The mean weight for each temperature group is indicated. Trout images were adapted from BioRender. **B.** Percentage of survival of trout infected by the parental strain IHNV_WT_ and the thermo-adapted recombinant variant rIHNV_25_, reared at 10°C, 15°C and 23°C. Mortalities were recorded every day for 26 days.

These three groups of fish, acclimated to different temperatures, were then subdivided into three groups of 25 individuals and infected by bath immersion with either the parental IHNV_WT_ or the recombinant thermo-adapted rIHNV_25_ viruses or mock infected with cell culture medium, as a negative control (Fig. 9B). CPM induced by IHNV_WT_ infection decreased as rearing temperature increased (final CPM of 44% at 10°C and 16% at 15°C) and no mortality was recorded at 23°C. Concerning rIHNV_25_ infection, as previously observed during the first trial, no mortality occurs at 10°C. The same observations were made at higher rearing temperatures (15°C and 23°C), indicating that rIHNV_25_ variant was completely attenuated in rainbow trout, regardless of the temperature, in contrast to the *in vitro* observations.

### Infection of zebrafish larvae by rIHNV_25_ and rIHNV_25_-mCherry and comparison of two rearing temperatures: 20°C versus 25°C

*In vivo* investigations of IHNV_25_ infectious process were next performed in zebrafish larvae using a well-established model for studying viral spreading and innate immune responses during infection with thermo-adapted IHNV variant [38, 39]. This fish lives at temperatures between 20°C and 28°C. The small size and optical transparency of zebrafish larvae enabled us to monitor the infection by fluorescence imaging at the whole-organism level using the traceable rIHNV_25_ variant expressing the red fluorescent protein mCherry (rIHNV_25_-mCherry) throughout the course of the infection. rIHNV_25_-mCherry was generated by reverse genetics as previously described [25]. The rIHNV_25_ genome was modified by the insertion, into the intergenic region between the M and G genes, of an expression cassette composed of the mCherry ORF, flanked by transcription gene start and gene end signals recognized by the viral L polymerase. Zebrafish larvae, at 3 days post fertilization (dpf), were infected by intravenous microinjections in the caudal vein with 2 nL of rIHNV_25_, rIHNV_25_-mCherry (60-80 PFU) or cell culture for mock-infected larvae, as previously reported [39]. They were then incubated at 20°C or 25°C. Appearance of clinical signs and mortality were assessed at 24, 48, 72, 96, 120 and 144 hours post infection (hpi) (Fig. 10).

**Fig. 10.**
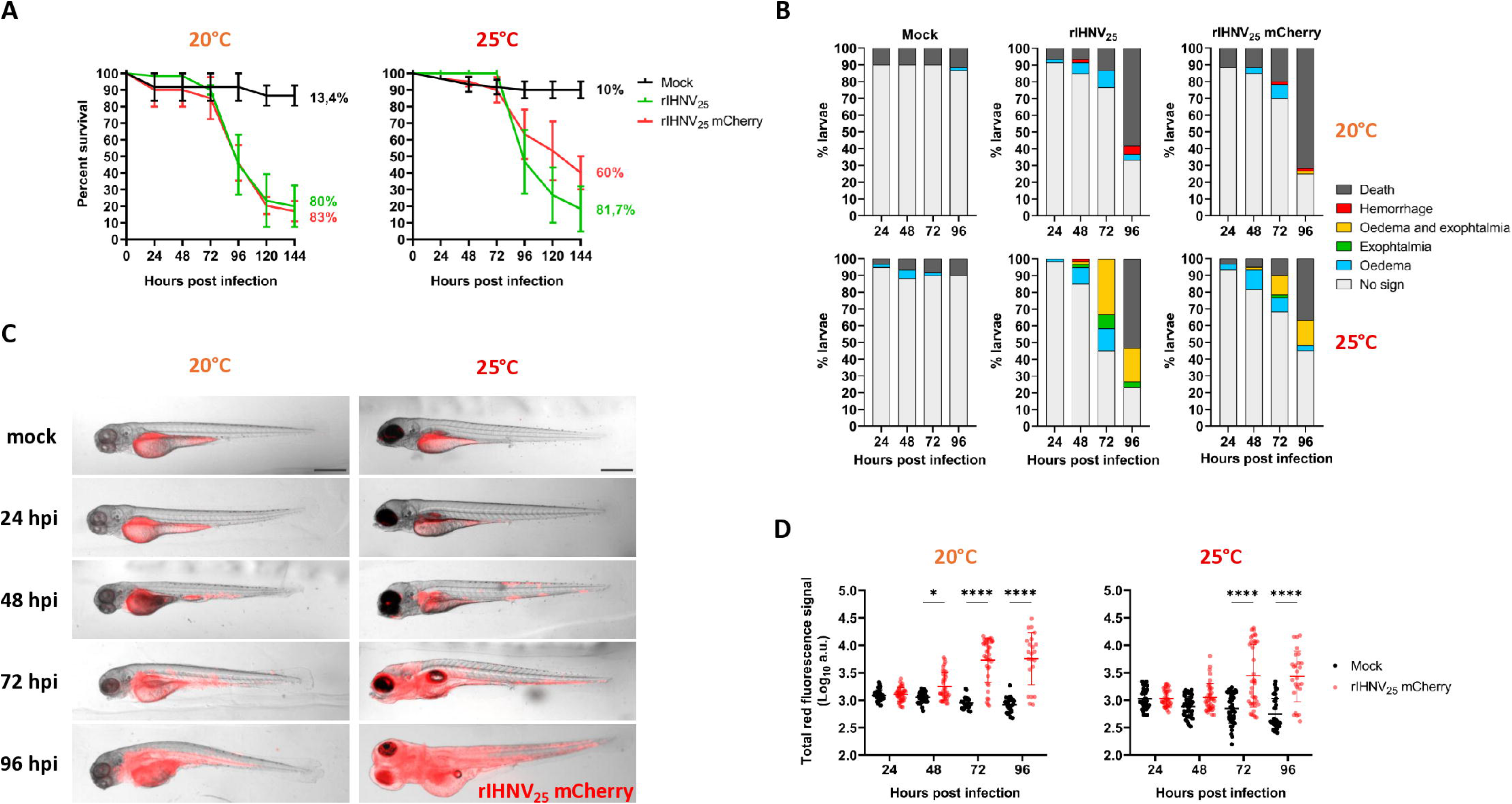
Zebrafish larvae infection model with rIHNV_25_ and rIHNV_25_-mCherry by microinjection at 20°C and 25°C. **A.** Wild-type AB and Nacre zebrafish larvae at 3 days post fertilization (dpf) were microinjected in the caudal vein with 2 nL of rIHNV_25_ or rIHNV_25_-mCherry (60-80 PFU), or with 2 nL of cell culture for mock-infected larvae and incubated at 20°C or 25°C (n = 20 per group). Mortality is presented as the percentage of survival recorded across three independent experiments. **B.** Infected larvae were observed daily post infection using a fluorescent stereomicroscope to detect any clinical signs. **C.** Examples of infection profiles observed in whole larvae infected with rIHNV_25_-mCherry at various times post infection. Scale bar: 200 μm. **D.** Quantification of fluorescence intensity in rIHNV_25_-mCherry-infected larvae using the COPAS system. Total red fluorescence signal of rIHNV_25_-mCherry-infected and mock-infected larvae, measured with COPAS system at various times post infection (mean of 20 larvae from two independent experiments, with standard errors). Statistically significant differences are indicated as follows: ****, p value < 0.0001 and *, p value < 0.05.

At both temperatures of 20°C and 25°C, the larvae were susceptible to rIHNV_25_, showing identical CPM of approximatively 80% at 144 hpi (Fig 10A). Clinical signs, such as oedema and hemorrhages, were observed as early as 48 hpi at 20°C (Fig. 10B). Similar CPM and clinical signs were also observed for the parental strain at 20°C (data not shown). Surprisingly, clinical signs induced by rIHNV_25_ were much more pronounced at 25°C, and a larger number of larvae exhibited oedema, exophthalmia, or both, as is often observed in rainbow trout infected with the IHNV (Fig. 10B and 10C) [12]. rIHNV_25_-mCherry induced clinical signs and mortality kinetics that were similar at 20°C but attenuated at 25°C (CPM of 60% for rIHNV_25_-mCherry versus 81.7% for rIHNV_25_), thereby confirming the attenuating effect of gene insertion as previously reported for several members of the *Mononegavirales* (Fig. 10A and 10B) [40–42]. To evaluate viral spreading over the same period, infected larvae were anesthetized to detect rIHNV_25_-mCherry signals by live fluorescence stereomicroscope imaging. In addition to the autofluorescence signals detected in the yolk sac of mock-infected fish at 20 and 25°C, red fluorescence signals specifically associated with the virus replication were observed as soon as 48 hpi in the infected fish (Fig 10C). mCherry fluorescence was initially detected in the immediate vicinity of the injection site (the caudal vein, below the urogenital orifice). The signal gradually spread distributing throughout the vascular network. At 96 hpi at 25°C, the larvae exhibited near-total red fluorescence. rIHNV_25_-mCherry infection was further quantified by flow cytometry using COPAS (complex object parametric analysis and sorting) technology. This continuous flow system allows the analysis of infected zebrafish larvae to detect fluorescent signals while maintaining their viability, as previously demonstrated for another fish rhabdovirus [42]. Larvae were analyzed daily between 24 to 96 hpi and the red fluorescence signals of mock-infected versus rIHNV_25_-mCherry infected larvae were analyzed. The technique proved sufficiently sensitive to detect mCherry fluorescence specifically associated with the virus replication as early as 48 hpi in the infected larvae, with intensities exceeding the background signal produced by the embryonic yolk in control larvae (Fig 10D). We also measured a continuous increase in mCherry fluorescence in infected larvae up to 96 hpi, confirming the high level of virus replication and spread within the larvae, as observed using a fluorescence stereomicroscope.

### rIHNV_25_ induces interferon and pro-inflammatory responses specific to the rearing temperatures

To investigate the effect of the temperature on the host response to rIHNV_25_ infection and its potential consequences for the severity of observed clinical signs, infected zebrafish larvae were incubated at 20°C or 25°C and were sampled between 0 to 56 hpi. The larvae were then processed to measure, by RT-qPCR, the expression of the rIHNV_25_ N gene (Fig. 11A and 11B) as well as that of two interferon genes (*ifnphi1* and *ifnphi3*), two interferon stimulated genes (ISGs: *viperin* also known as *vig1* or *rsad2* and *isg15*), and two proinflammatory markers of (*il1b* and *tnfa*) (Fig 11C and 11D).

**Fig. 11.**
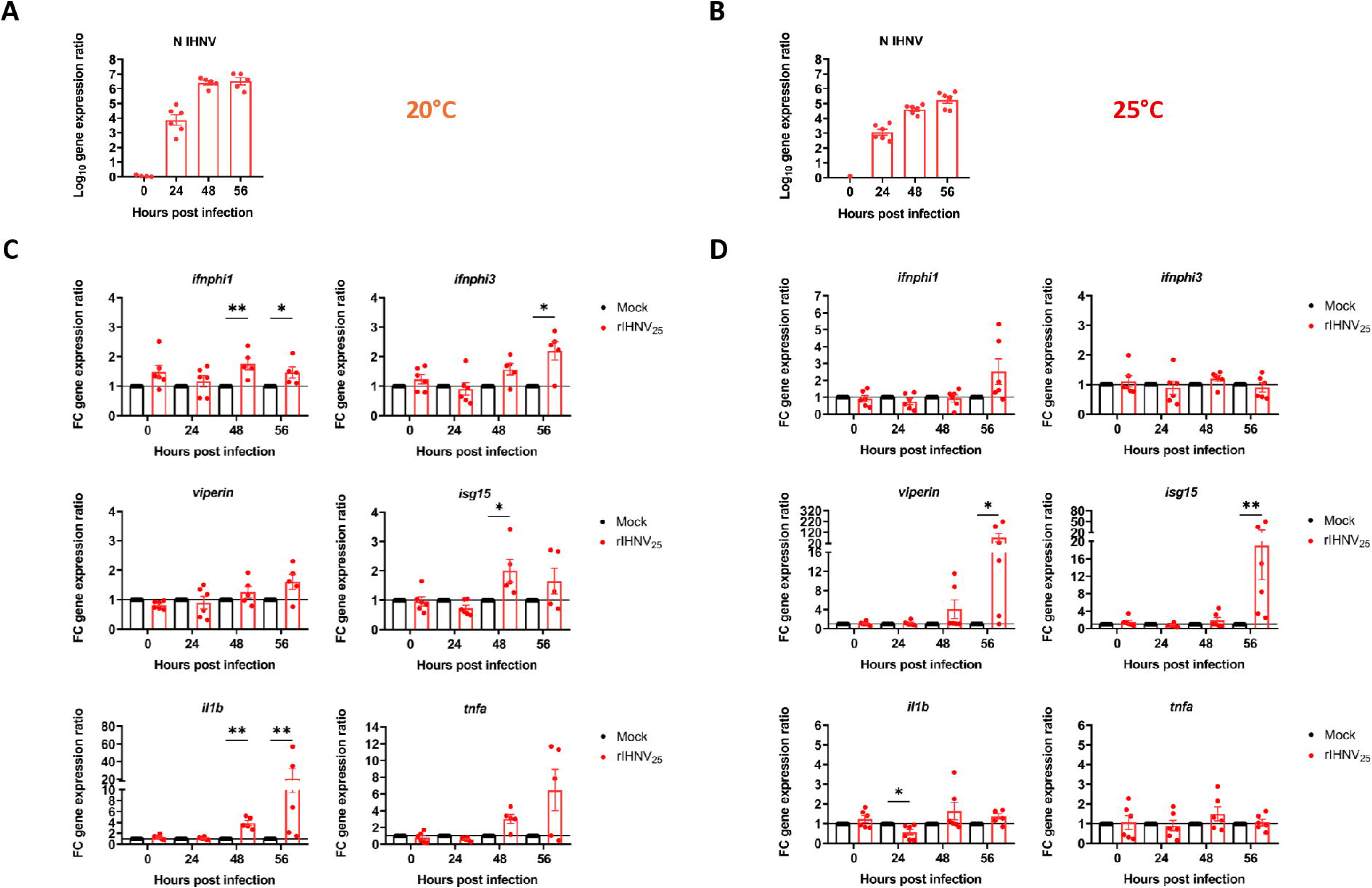
Analysis of the innate immune response in larvae infected by microinjection with rIHNV_25_, and comparison between 20°C and 25°C. Wild-type AB and Nacre zebrafish larvae at 3 dpf were microinjected in the caudal vein with either 2 nL rIHNV_25_ (60-80 PFU) or with 2 nL of cell culture medium for mock-infected larvae and reared at 20°C (A-C) or 25°C (B-D) (a total of n = 60 per group from two independent experiments). rIHNV_25_ replication in zebrafish larvae (A-B). Three groups of 5 larvae from 2 independent experiments were randomly harvested at different times post infection and gene expression was analyzed by RT-qPCR. Viral loads are expressed as the ratio of IHNV nucleoprotein N mRNA copy to that of the housekeeping gene *ef1a*. The expression of several genes involved in the antiviral response (*ifnphi1*, *ifnphi3*, *viperin*, *isg15*) and in the anti-inflammatory response (*il1b*, *tnfa*) were also measured (C-D) at 0, 24, 48 and 56 hpi. Each expression was normalized to the reference gene *ef1a*. Normalized expression values were standardized against their respective controls (Control fold change (FC) = 1, basal line). Graphs represent the mean of the fold changes ± standard error of the mean (SEM) of the biological replicates. Statistically significant differences are indicated as follows: **, p value < 0.01; *, p value < 0.05.

RT-qPCR measurements confirmed the active viral replication of rIHNV_25_ in zebrafish larvae, revealing a 6.5-log and 5.2-log increase in IHNV N-gene expression between 0 and 56 hpi, at 20°C and 25°C, respectively (Fig 11A and 11B). This result aligns with findings showing an increase in mCherry fluorescence in larvae infected with rIHNV_25_-mCherry, as measured by COPAS (Fig. 10D). The difference in rIHNV_25_ replication between the two temperatures, as measured by RT-qPCR, was also reflected in a decrease in mean mCherry fluorescence at 25°C compared with 20°C. Overall, these data demonstrate that rIHNV_25_ established a robust infection in zebrafish larvae at both 20°C and 25°C.

Infection induces a delayed and low-magnitude antiviral and inflammatory response at 20°C (Fig. 11C). Indeed, significant induction of interferon genes, ISGs, and proinflammatory markers occurs only between 48 to 56 hpi, just before the onset of mortality (Fig. 10A). In contrast, infection at 25°C induces strong ISG expression in the absence of interferon and inflammatory gene induction, although ISG induction was delayed and only occurred from 56 hpi onwards (Fig. 11D). Similar viral and immune gene expression profiles were observed when infections were performed at another permissive temperature of 21.5°C (Fig S1A and S1B), whereas no significant expression was detected during infections at a non-permissive temperature of 27°C (Fig S1C and S1D).

### rIHNV_25_ and rIHNV_25_-mCherry can infect zebrafish larvae by bath immersion at 25°C

To determine whether rIHNV_25_ had acquired the ability to “naturally” infect zebrafish larvae at 25°C, larvae at 3 dpf were infected by immersion in a bath containing 3 × 10^6^ PFU of rIHNV_25_ or rIHNV_25_-mCherry, or cell culture medium for mock-infected condition. The larvae were daily observed by live fluorescence stereomicroscopy to detect mCherry fluorescent foci, and mortality was recorded. Interestingly, some larvae were susceptible to rIHNV_25_ or rIHNV_25_-mCherry, showing an identical CPM of approximatively 35% at 168 hpi, with first death observed with in the rIHNV_25_ group (Fig. 12A). With the exception of one larva displaying mCherry signal in the trunk region, rIHNV_25_-mCherry entered the larvae via the caudal fin, as previously reported in rainbow trout [41]. Indeed, mCherry labelled primo-replication sites were mainly detected at the caudal fin lesions before detection in the head kidney, the target organ (Fig. 12B). Magnified views of the caudal fin illustrate the progressive spread of the virus and the increasing extent of the affected area over time (Fig. 12C).

**Fig. 12.**
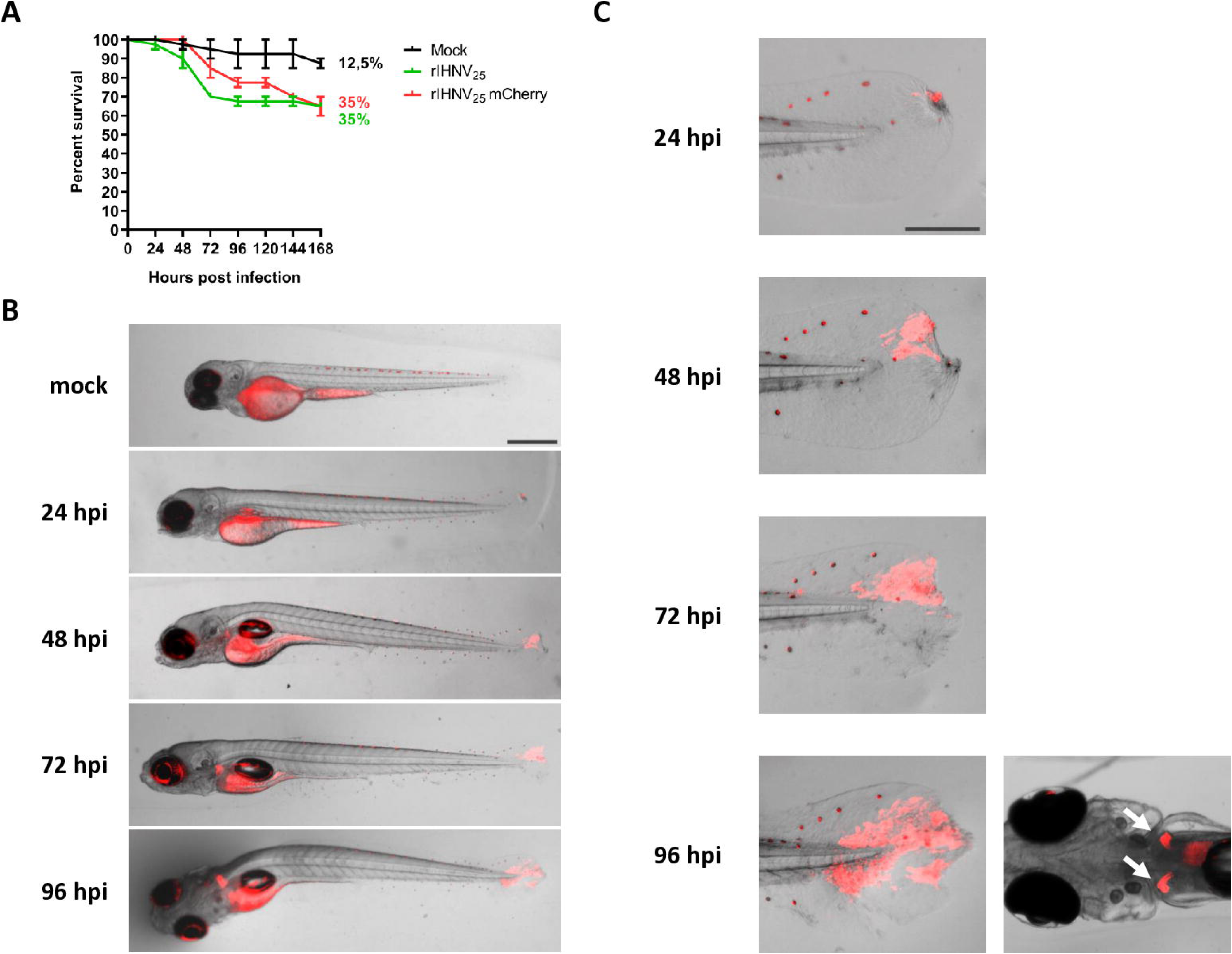
Zebrafish larvae infection with rIHNV_25_ and rIHNV_25_-mCherry by bath immersion at 25°C. **A.** Wild-type AB and Nacre zebrafish larvae at 3 dpf were infected by bath immersion with rIHNV_25_ or rIHNV_25_-mCherry (3 × 10^6^ PFU) or mock-infected with cell culture medium (20 larvae per group) and incubated at 25°C. Mortality is presented as the percentage of survival recorded in two independent experiments. **B.** Examples of rIHNV_25_-mCherry spread observed daily in the same infected larva with a fluorescent stereomicroscope. Scale bar: 200 μm. **C.** Magnified views of the caudal fin and head kidneys of the rIHNV_25_-mCherry infected larva from B. Scale bar: 100 μm.

## Discussion

In the early 1990s, experts began warning of the potentially exacerbating effects of climate change on the spread of infectious diseases, particularly those caused by terrestrial pathogens affecting humans and mammals. In the decades that followed, a growing body of work has emerged in this key research area [43]. However, these studies focus primarily on how climate change affects the geographic distribution, ecological dispersal, and epidemiology of infectious disease agents and their vectors in terrestrial environments. A recent meta-analysis revealed that 58% of human infectious diseases are aggravated by climate change [44]. Significant shifts in the geographic ranges of terrestrial host and vector species are cited as one of the most common ecological consequences of climate change regarding infectious diseases; global warming is thus linked to the range expansion of vectors such as mosquitoes, as well as host and reservoir species (birds and mammals) involved in outbreaks of dengue, chikungunya, Zika, and West Nile virus. In contrast, studies on the consequences of rising temperatures linked to climate change for aquatic animals, such as poikilothermic fish, have focused primarily on the host rather than on the fate of their pathogens. Indeed, it is well established that variations in ambient temperature have a considerable impact on the survival, growth and physiological parameters of fish [45] and can compromise their immunity, potentially increasing their susceptibility to disease [46]. However, from the pathogen’s perspective, what will be the intrinsic consequences of rising temperatures on its evolution and host range? In this study, we demonstrated that a Novirhabdovirus infecting cold-water fish could adapt to a gradual increase in temperature by acquiring nine mutations distributed across four of its six genes, a result verified by two independent experiments conducted with two different strains of IHNV. This adaptation extends its replication temperature range up to 25°C, while retaining its ability to replicate efficiently at low temperatures, like its parental strain.

The relationship between temperature fluctuations and viral evolution remains underexplored with few examples reported in the literature. One of the rare examples involved the use of the vesicular stomatitis virus (VSV) as a model [47]. Using an *in vitro* experimental evolution approach, the study showed that rhabdoviruses evolving under randomly fluctuating temperatures achieved lower fitness gains than those evolving under deterministically fluctuating temperatures. Although the study did not examine the effects of a gradual rise in temperature, which could mimic global warming scenarios, it suggests that RNA viruses, such as VSV, can rapidly acquire adaptive mutations distributed across the entire genome when evolving under conditions of fluctuating temperature conditions. Using reverse genetics, we previously demonstrated that it is possible to recover a recombinant IHNV in which the glycoprotein G gene has been replaced by that of the spring viraemia of carp virus (SVCV), a distantly-related rhabdovirus able to replicate at temperatures up to 28°C [34]. It is interesting to note that the recombinant virus rIHNV-G_SVCV_ replicates only within the temperature range of the parental IHNV strain (14 to 20°C), indicating that the glycoprotein is not sufficient to overcome this thermal restriction. In the present study, we demonstrated that three mutations within the glycoprotein G are essential for its correct conformation and/or targeting to the cell membrane at high temperature. It is noteworthy that the very first mutation appeared in the G protein as early as the 20°C stage (Fig. 2B). Surprisingly, these adaptive mutations in the G protein were observed in field strains recently isolated from rainbow trout in Europe, with some strains carrying two out of the three mutations (Table S1 and S2). This suggests that some strains might already present a certain advantage priming them to adapt rapidly to higher temperatures.

In another study focusing on VHSV, Kim and colleagues demonstrated that replacing the RNA polymerase L gene of a strain (MI03GL; genotype IVb), whose growth is sensitive to temperatures above 18 °C, with that of a divergent “exotic” strain (JF-09; genotype IVa) able to replicate at 23°C conferred a partial growth capacity at higher temperatures (>20°C) *in vitro* [35]. Although the study did not explore the contribution of the 31 amino acid substitutions observed between the L genes of these two genetically distant strains, it indicates that certain determinants of VHSV thermo-adaptation are located in the L gene and that other viral genes are also involved. In our study, we conclusively demonstrated that mutations in the L gene were essential, but that additional mutations in three other genes, namely N, M, and G, were also required, while mutations in the P gene could confer a replicative advantage at 25°C. Interestingly, sequencing intermediate variants isolated during the adaptation of IHNV to 24°C made it possible to determine the chronological order in which the amino acid substitutions appeared.

The mutations were mapped onto predictive structural models generated by the AlphaFold 3 algorithm. This provided a rational basis for structure-function analyses to identify domains in IHNV_25_ and IHNV_24_ proteins associated with adaptation to high temperatures. For N, P, G, and L proteins, the substitutions identified in the thermo-adapted strains map to distinct protein domains. Interestingly, for the M protein the substitution identified for both IHNV_25_ and IHNV_24_ involve the same residue position, M104 located in the globular domain of the protein which is substituted to an arginine (R) for IHNV_25_ and a threonine (T) for IHNV_24_. This result suggests that the site is particularly sensitive to temperature elevation and may be important for maintaining the protein’s stability or for protein-protein interactions. Using *in silico* analysis with Pythia [48], the effect of amino acid substitutions at position 104 on the stability of the matrix protein was evaluated and predicted to stabilize the protein (Fig S2). Most of the substitutions found in the G protein have already been identified in field isolates (5/6), while only one such conserved substitution was found for IHNV L protein (A1831T in the methyl transferase domain of IHNV_24_), this finding may reflect the limited number of complete IHNV genome sequences available in GenBank database. While locating potential thermo-adaptation mutational hotspots is not feasible using the only two strains available in this study, the comparative analysis of IHNV_25_ and IHNV_24_ substitutions maps indicates that there is no single mutational pathway to obtain thermo-adapted strains. This is likely because the adaptation at higher temperatures may require multiple stabilizing mutations and since the viral proteins are known to interact with each other. It is possible that compensatory mutations may have arisen during the process of thermo-adaptation, for example between N, P, and L proteins which form the viral replicase complex.

Interestingly, recent European isolates have already been reported as thermotolerant. Danish IHNV DK (2021), Italian IHNV 67-1 (2022), and French IHNV SA-22 (2022) strains replicated in cell culture at 15, 20°C but also at 22°C in contrast to historical IHNV strains [17, 18]. It should be noted that at a temperature of 25°C, these strains were no longer able to replicate *in vitro*. *In vivo*, in rainbow trout, the IHNV 67-1 isolate caused mortality at 12 and 19°C, whereas the IHNV DK isolate was virulent only at 12°C. Notably, although the virulence of the IHNV SA-22 isolate in rainbow trout remains under investigation, the virus caused 50% CPM in zebrafish larvae infected by bath immersion at a temperature of 23.5°C [17]. Comparison of the genomic sequences available for these strains with those of the *in vitro* thermo-adapted variants IHNV_24_ and IHNV_25_, described in the present study, reveals that the only shared amino-acid substitutions were identified in the glycoprotein: H98Y, common to IHNV_25_, SA-22 and 67-1 and A57V common to IHNV_24_ and 67-1 (Table S1 and S2). Interestingly the A57V substitution was the first to appear during the thermo-adaptation process of IHNV_24_ at 20°C (Fig. 2B). This highlights the importance of acquiring specific mutations in the glycoprotein of IHNV to gain a thermotolerant phenotype. These two mutations warrant further study to clearly demonstrate their role in the acquisition of this phenotype.

It is interesting to note that the rIHNV_25_ variant, while now replicating at 25°C, has retained efficient replication at low temperatures, like its parental strain, in the cyprinid-derived EPC cell line (Fig. 5B), thereby expanding its replication temperature range. In contrast, replication of rIHNV_25_ was delayed in the salmonid-derived cell line compared to that of its parental strain, except at 23°C, a temperature non-permissive for the parental strain (Fig. 5B). In rainbow trout, rIHNV_25_ variant was completely attenuated, regardless of temperature (Fig. 9B). It is possible that, among the 11 mutations identified in its genome, some are linked to adaptation to cell culture, a likely consequence of repeated passages in a cyprinid cell line, rather than to thermo-adaptation itself. This type of mutation adapting to a new host could underlie the attenuation of infection and/or viral replication in the natural host. However, it cannot be ruled out that mutations related to the thermo-adaptation also have an attenuating effect on rIHNV_25_ in trout, even though the fish were reared at higher temperature (23°C). rIHNV_25_ was also tested in zebrafish larvae, a warm-water species, to determine the potential for crossing the species barrier associated with the thermo-adaptation. This fish lives at temperatures between 20°C and 28°C and is not naturally infected with IHNV. At 20°C and 25°C, high mortality rates were observed in larvae microinjected with rIHNV_25_, and rIHNV_25_-mCherry into the caudal vein. Infection was associated with severe symptoms identical to those observed in trout (hemorrhages, oedema, and exophthalmia). At both temperatures, mortality was preceded by a massive viral replication and a delayed and inefficient interferon and inflammatory response, as previously reported [39, 49]. Surprisingly, IHNV_25_ was also capable of infecting larvae by bath immersion at 25°C, a non-permissive temperature for IHNV field strains *in vitro* [17, 18], and entering via the caudal fin, as reported in trout [41], thus demonstrating its ability to cross the species barrier through its adaptation to higher temperatures.

This study examined a major threat to global aquaculture by characterizing novirhabdoviruses that adapt to rising temperatures, as well as the consequences for the fish hosts. It sheds light on the molecular and evolutionary processes that underlie the emergence of viral strains adapted to higher temperatures, as well as their potential virulence towards other fish species. The identification of viral markers of thermo-adaptation will be of great importance for the epidemiological surveillance of IHNV, a virus regulated in EU that must be notified to the WOAH. These markers will enable to update diagnostic and predictive characterization tools and facilitate the detection of potential emerging strains. This study provides an innovative framework that can be extended to the study of the impact of climate change on other rhabdoviruses (610 species in 63 genera) [50], including ectothermic insect-vectored rhabdoviruses that infect terrestrial crop plants or farmed mammals (e.g. Bovine ephemeral fever virus (BEFV) transmitted by mosquitoes) thus providing valuable insights for other essential food-producing sectors. More broadly, this approach can be extended to all viruses infecting poikilothermic organisms, including all viral pathogens infecting cold-water fish.

## Materials and methods

### Ethics Statements

All animal studies were conducted in strict compliance with European guidelines and recommendations regarding animal experimentation and welfare (European Union Directive 2010/63). All experimental procedures involving animals were approved by the local ethics committee on animal experimentation (Comité d’éthique appliqué à l’expérimentation animale pour le centre de recherches INRAE de Jouy-en-Josas; COMETHEA no. 45) and authorized by the Ministère de l’Éducation nationale, de l’Enseignement supérieur et de la Recherche under authorization numbers: APAFIS#29801-2021021110262075 v2 and APAFIS#46003-2023112017042104, as well as for the fish facilities (authorization number D-78-322-720). Lethal experimental infection with IHNV typically results in acute disease characterized by anemia, exophthalmia and petechial hemorrhages. Consequently, fish were monitored at least two times daily to detect clinical signs and assess survival. Upon the onset of characteristic symptoms, the animals were humanely euthanized by immersion in a bath containing a lethal dose of tricaine (MS-222 Sigma at 300 mg/L).

### Cells and viruses

Parental strains and thermo-adapted variants of IHNV were propagated in EPC (Epithelioma Papulosum Cyprini from fathead minnow, *Pimephales promelas*) cell monolayers at 15°C, as previously described [32, 51]. Virus titers were determined by plaque assay on EPC cells under an Avicel overlay (1.2% Avicel RC581 (FMC Biopolymer) in Glasgow’s modified Eagle’s medium at pH 7.6 with Tris-HCl 23.75 mM and HCO_3_Na 4 mM and supplemented with 2% fetal bovine serum and 2 mM L-glutamine). At 3 to 5 days post infection, Avicel containing medium was removed, and the cell monolayers were fixed and stained with crystal violet. Recombinant vaccinia virus expressing the T7 RNA polymerase, vTF7-3 [52], was kindly provided by B. Moss (National Institutes of Health, Bethesda, MD). For multiple-step growth kinetics, EPC and rainbow trout (*Oncorhynchus mykiss*) RTgill-W1 cells [53] (grown in Leibovitz’s L-15 medium and kindly provided by C. Tafalla (National Institute of Agricultural and Food Research and Technology, Madrid, Spain)) were infected at an MOI of 0.01 PFU per cell. After one hour of adsorption, the inocula were removed and the cell monolayers were washed twice before being overlaid with 1 mL of fresh medium and incubated at 10°C, 15°C, 20°C, 23°C or 25°C. Supernatant aliquots (0.1 mL out of a total medium volume of 1 mL per well) were collected at various times post infection and replaced by an equivalent volume of fresh medium. The samples were flash-frozen and subsequently analyzed by plaque assay.

### Adaptation of the virus to a higher incubation temperature and whole genome sequencing

A recombinant IHNV, derived from the French isolate 32-87 and expressing the mCherry fluorescent protein (rIHNV-mCherry [25]), was progressively adapted to 24°C by infecting EPC cells at increasing temperatures (as shown in Figure 1). All rIHNV-mCherry intermediates, obtained at the different temperature plateaus, were subjected to high-throughput sequencing to determine the complete consensus sequences of their genome. Sample processing procedures, as well as sequencing and bioinformatic analyses, are described in detail in Zamperin et al. 2019 [54].

### Prediction of IHNV CAN67 and 32-87 protein structures

To predict the structures of the parental strain proteins used in the study, the amino acid sequences of the N, P, M, G, and L proteins were submitted to AlphaFold 3 server (https://alphafoldserver.com) [55]. The NV protein was not included in the analysis as no substitutions were found for both IHNV_25_ and IHNV_24_. Structural modeling was performed for both parental strains except for P, where only the CAN67 structure was predicted (parental strain of IHNV_25_) because no substitutions were found for IHNV_24_ P. IHNV P and G proteins were predicted as a dimer or a trimer, respectively, based on available structural data for other rhabdoviral proteins. Domain mapping for IHNV N and P [26], M [27], G [30], and L [29] proteins were based on the known protein domain organization found in published studies on vesicular stomatitis virus (VSV) and rabies virus (RABV) protein structures and corresponding protein alignments. Structural representations and protein domain labeling were performed using ChimeraX [56].

### Construction of a cDNA clone of a thermo-adapted variant of IHNV and recovery of recombinant viruses

A full-length cDNA copy of the RNA genome of the thermo-adapted IHNV variant was amplified as five overlapping cDNA fragments by reverse transcription-PCR (RT-PCR) and were fully sequenced using specific primers, as previously described [32]. Based on this sequence, a full-length cDNA copy was synthetized *in vitro* and cloned in a pUC57 plasmid (Genscript). The final cDNA construct, pIHNV_25_, consists of an antigenomic cDNA copy of the RNA genome of the thermo-adapted IHNV variant, cloned under the control of a T7 promoter and upstream a hepatitis delta virus-derived ribozyme sequence and a T7 terminator sequence. This construct was subsequently modified to introduce an expression cassette encoding the red fluorescent protein mCherry. The expression cassette was designed as previously described [57] and inserted in the intergenic region between the M and G genes of pIHNV_25_, leading to pIHNV_25_-mCherry (Table 1).

Chimeric constructs between the parental strain and the thermo-adapted variant were obtained by replacing homologous fragments using unique restriction enzyme sites as illustrated in Figure 6. In some cases, a fragment was amplified by PCR and subcloned in a pJet1.2 vector (Thermo Fischer Scientific) before serving as a template for site-directed mutagenesis using specific primers (Table 1). All final constructs were sequenced.

Recovery of rIHNV was performed as previously described [32]. Approximately 5 × 10^6^ EPC cells per well were grown in six-well plates and infected with the recombinant vaccinia virus vTF7-3, expressing the T7 RNA polymerase, at an MOI of 5. After 1 h of adsorption at 37°C, the cells were washed twice and transfected with a plasmid mixture containing 0.25 μg of pT7-N, 0.2 μg of pT7-P, 0.2 μg of pT7-L and 1 μg of the full-length cDNA construct, using Lipofectamine 2000 (Thermo Fisher Scientific) according to the supplier’s instructions. The cells were incubated for 6 h at 37°C; the transfection mixture was replaced with fresh medium, and the cells were then incubated at 15°C for 7 days. Subsequently, the cells were harvested by scraping the plates with a rubber policeman and subjected to two of freeze-thaw cycles. The cell culture supernatant (called P0 supernatant) was clarified by centrifugation at 6,000 × *g* for 10 min in a microcentrifuge at 4°C and used to inoculate fresh EPC cell monolayers in 24-well plates at 15°C.

### Plasmid constructs, transfection, indirect immunofluorescence analyses on fixed and live cells and Western-blot analyses

Plasmid vectors expressing glycoproteins derived from the parental strain and the thermo-adapted variant were constructed by amplifying the G genes of each virus using specific primers (Table 1) and cloning them into a pcDNA1.1/Amp. EPC cells were seeded in 6-well plates at a concentration of 5 × 10^6^ cells per well, 24 h prior to transfection by electroporation (Nucleofector technology; Lonza). The cells were trypsinized and resuspended in 100 μL of solution T. They were then mixed with 1 μg of plasmid DNA, electroporated using program T-020, distributed onto two cell culture slides (Ibidi), and incubated at 15°C or 20°C. At 48 h post transfection, cells were fixed with a mixture of alcohol and acetone [1:1 (v/v)] at −20°C for 15 min. Antigen detection was performed by incubation with a mouse monoclonal antibody (mAb) 19B7 (dilution 1:1,000) against IHNV G protein diluted in PBS1x containing 0.05% of Tween 20 for 45 min at room temperature (RT). The cells were then washed three times, incubated with Alexa Fluor 594-conjugated anti-mouse immunoglobulins antibodies (dilution 1:1,000; Invitrogen), and nuclei were stained with Hoechst (dilution 1:1,000; Thermo scientific) for 45 min at RT. Three washes were performed, and fluoromount G (SouthernBiotech) was added prior to imaging. For live-cell imaging, transfected cell monolayers were directly incubated with a mouse mAb 6H9 (dilution 1:1,000) against IHNV G protein diluted in GMEM cell culture medium supplemented with 10% FBS for 45 min at RT. After 3 washes with the same medium, the cells were incubated with Alexa Fluor 594-conjugated anti-mouse immunoglobulin antibodies (dilution 1:1,000; Invitrogen) and nuclei were stained with Hoechst (dilution 1:1,000; Thermo scientific), for 45 min at RT. Three washes were performed prior to imaging. The cell monolayers were then observed using an Axio Observer.Z1 inverted fluorescence microscope (Zeiss) equipped with a CoolSNAP HQ2 CCD camera (Photometrics Roper Scientific). Microscopy images were acquired using Zen 2.3 software (Zeiss).

For protein detection, transfected cells cultured in 6-well plates were lysed at 48 h post transfection using 150 μL of lysis buffer consisting of 150 mM NaCl, 50 mM Tris-HCl (pH 8.5), 20 mM EDTA, 1% deoxycholic acid sodium salt (DOC), and 1% Triton X-100. Cell lysates were homogenized using QIAshredder columns (Qiagen). Aliquots of 100 μL were then incubated with 25 μL of cracking buffer composed of 42% glycerol, 21% β-mercaptoethanol, 250 mM Tris (pH 7), 12.5 % SDS, and bromophenol blue for 5 min in boiling water. The samples were subsequently separated by electrophoresis on a 4 to 12% gradient Bis-Tris polyacrylamide gel (NuPage Novex; Invitrogen) followed by Western blot analysis. After electro-transfer onto a polyvinylidene difluoride (PVDF) membrane (Immobilon-P; Millipore), proteins were detected using a mouse mAb 19B7 (dilution 1:2,000), directed against IHNV G protein, and a mouse anti-alpha tubulin antibody (Sigma) diluted to 1:3,000 for loading controls. Immunodetected antigens were visualized with a peroxidase (HRP)-conjugated goat anti-mouse antibody diluted to 1/5,000 (SeraCare) and an enhanced chemiluminescence (ECL) detection system (Pierce). Antigen detection was finally performed using Amersham ImageQuant 800 Western Blot Imaging System (Cytiva). The expression ration of IHNV G protein to alpha-tubulin were calculated for each condition.

### Experimental infection of rainbow trout

Virus-free juvenile INRAE AUT*SYN or “maigre” line of reference (n = 25 or 50 per group) of rainbow trout (*Oncorhynchus mykiss*) (mean weight of 0.54 g to 1.27 g) were reared at 10°C or gradually acclimated to 15°C and 23°C starting from 10°C (see acclimation process in Figure 9). Trout were infected by immersion in thermoregulated tanks filled with 3 liters of freshwater for 2 hours with the parental strain IHNV_WT_ (IHNV CAN67), the thermo-adapted variant IHNV_25_ or the thermo-adapted recombinant virus rIHNV_25_ (final titer, 5 × 10^4^ PFU/mL) at 10°C, 15°C or 23°C. Tanks were then filled to a volume of 20 liters with freshwater (dechlorinated tap water), and the water flow was activated (open-circuit system, 1 renewal per hour). Tank temperature and oxygen concentration was monitored continuously (1 measurement every 10 minutes, @HACH Lange LDO sensors). Controls were fish mock infected with cell culture medium under the same conditions. Clinical signs and mortalities were recorded twice a day over a period of 26 to 28 days. For the group infected by rIHNV_25_ variant, surviving trout were challenged by immersion at day 28 with the IHNV_WT_ as described above (final titer, 5 × 10^4^ PFU/mL). Mortality was recorded daily for 29 days.

### IHNV infection of zebrafish larvae

Zebrafish (*Danio rerio*) were maintained at 28°C on a 14 h light/10 h dark cycle. Fish were housed in the animal facility of our laboratory which was built according to the respective local animal welfare standards [58]. Wild-type AB and Nacre zebrafish larvae [59] at 3 days post fertilization (dpf) were infected by microinjection under anesthesia with 250 mg/L tricaïne solution (MS-222, Sigma), as previously described [42]. Larvae were microinjected in the caudal vein with 2 nL of IHNV_WT_, rIHNV_25_ or rIHNV_25_-mCherry (60-80 PFU) or with 2 nL of cell culture for mock-infected larvae with a glass microneedle using a mechanical micromanipulator (M-152, Narishige), and a Femtojet microinjector (Eppendorf). Subsequently, the larvae were maintained in 1× E3 solution supplemented with 0.0003% methylene blue (Sigma) at a temperature of 20°C or 25°C. Infection by bath immersion was also tested on 3 dpf zebrafish larvae with rIHNV_25_ or rIHNV_25_-mCherry (3 × 10^6^ PFU) or mock-infected with cell culture medium (20 larvae per group). Viral inocula were diluted in a sterile 1× E3 solution containing 0.0003% methylene blue in a final volume of 200 µL and incubated at 25°C. Larvae were observed daily using an MZ10F fluorescent stereomicroscope (Leica Microsystems) to detect mCherry fluorescent foci, and mortality was recorded. Experimental infections by bath immersion or microinjection were repeated two or three times, respectively.

The COPAS XL large-particle sorter (Union Biometrica) was used to analyze zebrafish larvae that were either infected with the rIHNV_25_-mCherry virus or mock-infected at 0, 24, 48, 72 and 96 h post infection and at the two temperatures tested, as previously described [42]. Data were analyzed using the integrated mCherry fluorescence value for the entire body of the larvae.

RT-qPCR was used to evaluate the expression levels of immune and inflammatory genes of zebrafish larvae infected or not with rIHNV_25_, and incubated at different temperatures (20°C, 21.5°C, 25°C or 27°C). Viral replication was confirmed by measuring the relative expression of the gene encoding the IHNV N protein as previously described [39]. Briefly, rIHNV_25_-infected or mock-infected larvae were harvested and pooled in batches of 5 individuals at 0, 24, 48 and 56 hpi (three batches per time point and temperature). Larvae were homogenized in 350 µL of RLT buffer (Qiagen), and samples were stored at -80°C. RNA was extracted using the RNeasy Mini Kit (Qiagen), including a DNase I treatment step (Qiagen), in accordance with the manufacturer’s protocol. Total RNA quantity in each sample was determined using a NanoDrop 2000c spectrophotometer (Thermo Scientific), and cDNA was synthesized from total RNA using PrimeScript RT Master Mix (TaKaRa) for subsequent qPCR analysis. Finally, qPCR reactions were performed in a 25 µL reaction volume containing 12.5 µL of iTaq Universal SYBR Green Supermix (Bio-Rad), 8.5 µL of RNase-free water, 1 µL of each specific primer (10 µM), and 2 µL of cDNA template (diluted 1:4 in RNase-free water). All reactions were performed in duplicate (technical replicates) on a CFX Duet Real-Time PCR system (Bio-Rad), with an initial denaturation step (95°C, 30 s) followed by 40 cycles comprising a denaturation step (95°C, 5 s) and an annealing-elongation step (60°C, 30 s). Relative expression levels of the target genes were normalized using the 2^−ΔΔCT^ method [60], with the *eukaryotic translation elongation factor 1 alpha 1 like 2 (ef1a)* serving as the reference gene.

### Statistical analysis

Plaque size shown in Figure 4B represent means ± standard errors (SE) measured from approximately 15 plaques. One-way ANOVA comparisons were performed using analysis of variance (Kruskal-Wallis test) and Dunn’s multiple comparison test, with the p-values < 0.05 considered significant.

Kinetics of survival for groups infected with the parental strain, the thermo-adapted variant and the thermo-adapted recombinant virus were analyzed by Kaplan–Meier estimation of survival curves, followed by comparison of the survival curves by Log-rank test (Mantel–Cox) for significant differences (p values <0.05 considered significant).

Results of the individual red fluorescence in each larva, detected using the COPAS system, as well as gene expression results analyzed by RT-qPCR, were plotted as the mean ± standard error of the mean (SEM), and significant differences were determined using Kruskal-Wallis test and displayed as ****, p value < 0.0001; ***, p value < 0.001; **, p value < 0.01; and *, p value < 0.05. All analyses were performed using GraphPad Prism software version 9.0.

## Acknowledgments

We thank Monique LeBerre for the gift of IHNV_25_ and Michel Brémont for the initial genome sequence of IHNV_25_. We are grateful to members of the fish facilities, Magalie Bouvet, Pénélope Simon, Damien Degallaix and Kilian Cella, for taking care of experimental fish (IERP-UE907, Jouy-en-Josas Research Center, France doi.org/10.15454/1.5572427140471238E12) which belongs to the National Distributed Research Infrastructure for the Control of Animal and Zoonotic Emerging Infectious Diseases through In Vivo Investigation (EMERG’IN DOI: doi.org/10.15454/90CK-Y371). We thank Dr Carolina Tafalla for the gift of RTgill-W1 cells.

## Author contributions

Author contributions following the CRediT taxonomy (https://casrai.org/credit/) are as follows: Conceptualization: SB, JKM, CL; Formal analysis: SB, JKM, CF; Funding acquisition: SB, JKM, CL, YB, TM; Investigation: CF, EG, MM, JB, AL, MA, EM, MF, LL, YB, TM, DR, CL, JKM, SB; Methodology: CF, CL, MM, DM; Project administration: SB, JKM, CL, YB, TM; Supervision: SB, JKM, CL, TM; Visualization: CL, JKM, SB; Writing – original draft: SB, JKM, CF; Writing – review & editing: CF, EG, MM, JB, AL, MA, EM, MF, LL, YB, TM, DR, CL, JKM, SB.

## Supplemental figures

**Fig. S1. Analysis of the innate immune response in larvae infected by microinjection with rIHNV_25_ at 21.5°C and 27°C.**

Wild-type AB and Nacre zebrafish larvae at 3 dpf were infected with rIHNV_25_ by microinjection in the caudal vein or mock-infected with cell culture medium, as described in Figure 10, and reared at 21.5°C (A-B) or 27°C (C-D) (n = 20 per group from two independent experiments (A-B) or a single experiment (C-D). rIHNV_25_ replication in zebrafish larvae (A-C). Three groups of 5 larvae for each experiment were randomly harvested at different times post infection and gene expression was analyzed by RT-qPCR. Viral loads are expressed as the ratio of IHNV nucleoprotein N mRNA copy to that of the housekeeping gene *ef1a*. The expression of genes involved in the antiviral response (*ifnphi1*, *ifnphi3*, *viperin*, *isg15*) and in the anti-inflammatory response (*il1b*, *tnfa*) were also measured (B-D) at 0, 24, 48 and 56 hpi. Each expression was normalized to the reference gene *ef1a*. Normalized expression values were standardized against their respective controls (Control fold change (FC) = 1, basal line). The graphs represent the mean of the fold changes ± SEM of the biological replicates. Statistically significant differences are indicated as follows: ***, p value < 0.001; **, p value < 0.01; *, p value < 0.05.

**Fig. S2. Estimation of the effects of M104R and M104T substitutions on M protein stability using Pythia.**

The AlphaFold-predicted structures of CAN67 (**A**) and 32-87 (**B**) M proteins (shown in box insets) were used as input for Pythia protein stability analysis (https://pythiastudio.wulab.xyz/tools/pythia). For each position and amino acid substitution, Pythia calculates an energy score that provides an estimation of the stability change after such substitution that corresponds to the difference in the Gibbs free energy of unfolding, ΔΔG. Pythia uses a scoring system where a negative score indicates that the mutation is stabilizing (blue), while a positive value indicates that the mutation is destabilizing (red). The heatmaps are centered on residue position 104 of each M protein.

---

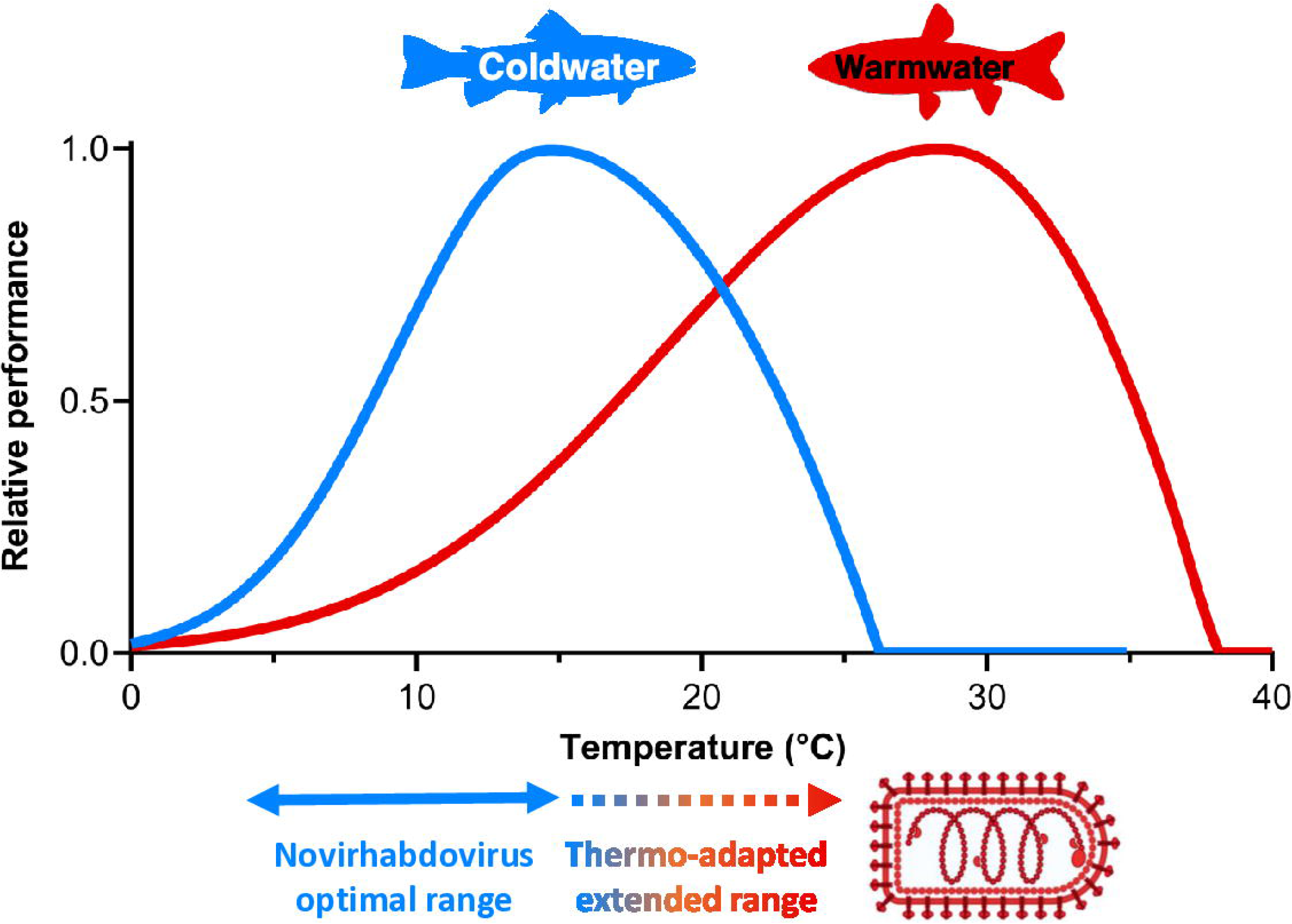

## References

1. Ripple, W.J., et al., The 2023 state of the climate report: Entering uncharted territory. BioScience, 2023. 73(12): p. 841–850.

2. FAO, The State of World Fisheries and Aquaculture 2022. Towards Blue Transformation. Rome, FAO. 2022.

3. Feist, S.W., et al., The aquatic animal pandemic crisis. Rev Sci Tech, 2019. 38(2): p. 437–457.

4. Quan, J., et al., Proteome analysis of rainbow trout (Oncorhynchus mykiss) liver responses to chronic heat stress using DIA/SWATH. J Proteomics, 2021. 233: p. 104079.

5. Gobler, C.J., et al., Temperature, Acidification, and Food Supply Interact to Negatively Affect the Growth and Survival of the Forage Fish, Menidia beryllina (Inland Silverside), and Cyprinodon variegatus (Sheepshead Minnow). Frontiers in Marine Science, 2018. 5.

6. Paez, D.J., et al., Temperature Variation and Host Immunity Regulate Viral Persistence in a Salmonid Host. Pathogens, 2021. 10(7).

7. Kemp, L., et al., Climate Endgame: Exploring catastrophic climate change scenarios. Proc Natl Acad Sci U S A, 2022. 119(34): p. e2108146119.

8. Moss, R.H., et al., The next generation of scenarios for climate change research and assessment. Nature, 2010. 463(7282): p. 747–56.

9. Janhunen, M., et al., Thermal sensitivity of growth indicates heritable variation in 1-year-old rainbow trout (Oncorhynchus mykiss). Genet Sel Evol, 2016. 48(1): p. 94.

10. Abbadi, M., et al., Increased virulence of Italian infectious hematopoietic necrosis virus (IHNV) associated with the emergence of new strains. Virus Evol, 2021. 7(2): p. veab056.

11. Kantala, T., et al., The first detection of infectious hematopoietic necrosis virus (IHNV) genogroup U in western Europe in the first IHNV outbreak in Finland. Res Vet Sci, 2026. 209: p. 106271.

12. Biacchesi, S. and M. Brémont, Vaccination against viral hemorrhagic septicemia and infectious hematopoietic necrosis, in Fish Vaccination, R. Gudding, A. Lillehaug, and Ø. Evensen, Editors. 2014, John Wiley & Sons, Ltd, : Chichester, UK. p. 334–340.

13. Kurath, G. and J.C. Leong, Characterization of infectious hematopoietic necrosis virus mRNA species reveals a nonvirion rhabdovirus protein. J Virol, 1985. 53(2): p. 462–8.

14. Niner, M.D., et al., Genomic and immunogenic changes of Piscine novirhabdovirus (Viral Hemorrhagic Septicemia Virus) over its evolutionary history in the Laurentian Great Lakes. PLoS One, 2021. 16(5): p. e0232923.

15. Troyer, R.M. and G. Kurath, Molecular epidemiology of infectious hematopoietic necrosis virus reveals complex virus traffic and evolution within southern Idaho aquaculture. Dis Aquat Organ, 2003. 55(3): p. 175–85.

16. Einer-Jensen, K., et al., Evolution of the fish rhabdovirus viral haemorrhagic septicaemia virus. J Gen Virol, 2004. 85(Pt 5): p. 1167–79.

17. Gourhannic, E., et al., Epidemic Risks associated to high-temperature replicating infectious hematopoietic necrosis virus (IHNV) field isolates: from molecular characterization to host pathogen interactions. Submitted, 2026.

18. Marsella, A., et al., Temperature impact on replication and virulence of European infectious hematopoietic necrosis viruses. Aquaculture, 2025. 609: p. 742786.

19. de Kinkelin, P., M. Bearzotti-Le Berre, and J. Bernard, Viral hemorrhagic septicemia of rainbow trout: selection of a thermoresistant virus variant and comparison of polypeptide synthesis with the wild-type virus strain. J Virol, 1980. 36(3): p. 652–8.

20. Cascarano, M.C., et al., Mediterranean Aquaculture in a Changing Climate: Temperature Effects on Pathogens and Diseases of Three Farmed Fish Species. Pathogens, 2021. 10(9).

21. Fay, R.L., et al., Experimental Evolution of West Nile Virus at Higher Temperatures Facilitates Broad Adaptation and Increased Genetic Diversity. Viruses, 2021. 13(10).

22. Tanaka, S., H. Aoki, and T. Nakai, Pathogenicity of the Nodavirus Detected from Diseased Sevenband Grouper Epinephelus septemfasciatus. Fish Pathology, 1998. 33(1): p. 31–36.

23. Combe, M., et al., Impact of Global Warming on the Severity of Viral Diseases: A Potentially Alarming Threat to Sustainable Aquaculture Worldwide. Microorganisms, 2023. 11(4).

24. Amend, D.F., W.T. Yasutake, and R.W. Mead, A Hematopoietic Virus Disease of Rainbow Trout and Sockeye Salmon. Transactions of the American Fisheries Society, 1969. 98(4): p. 796–804.

25. Biacchesi, S., et al., Limited interference at the early stage of infection between two recombinant novirhabdoviruses: viral hemorrhagic septicemia virus and infectious hematopoietic necrosis virus. J Virol, 2010. 84(19): p. 10038–50.

26. Ribeiro, E.A., Jr., et al., Dimerization of Rabies Virus Phosphoprotein and Phosphorylation of Its Nucleoprotein Enhance Their Binding Affinity. Viruses, 2024. 16(11).

27. Jenni, S., et al., Visualizing molecular interactions that determine assembly of a bullet-shaped vesicular stomatitis virus particle. Nat Commun, 2022. 13(1): p. 4802.

28. Roche, S., et al., Structure of the prefusion form of the vesicular stomatitis virus glycoprotein G. Science, 2007. 315(5813): p. 843-8.

29. Liang, B., et al., Structure of the L Protein of Vesicular Stomatitis Virus from Electron Cryomicroscopy. Cell, 2015. 162(2): p. 314–327.

30. Vollmer, B., et al., The prefusion structure of herpes simplex virus glycoprotein B. Sci Adv, 2020. 6(39).

31. Kim, C.H., J.R. Winton, and J.C. Leong, Neutralization-resistant variants of infectious hematopoietic necrosis virus have altered virulence and tissue tropism. J Virol, 1994. 68(12): p. 8447–53.

32. Biacchesi, S., et al., Recovery of NV knockout infectious hematopoietic necrosis virus expressing foreign genes. J Virol, 2000. 74(23): p. 11247–53.

33. Brudeseth, B.E., H.F. Skall, and O. Evensen, Differences in virulence of marine and freshwater isolates of viral hemorrhagic septicemia virus in vivo correlate with in vitro ability to infect gill epithelial cells and macrophages of rainbow trout (Oncorhynchus mykiss). J Virol, 2008. 82(21): p. 10359–65.

34. Biacchesi, S., et al., Heterologous exchanges of the glycoprotein and the matrix protein in a Novirhabdovirus. J Virol, 2002. 76(6): p. 2881–9.

35. Kim, S.H., et al., Interchange of L polymerase protein between two strains of viral hemorrhagic septicemia virus (VHSV) genotype IV alters temperature sensitivities in vitro. Virus Res, 2015. 195: p. 203–6.

36. Dixon, P., et al., Epidemiological characteristics of infectious hematopoietic necrosis virus (IHNV): a review. Vet Res, 2016. 47(1): p. 63.

37. Lapatra, S.E., et al., Size-Related Susceptibility of Salmonids to Two Strains of Infectious Hematopoietic Necrosis Virus. Transactions of the American Fisheries Society, 1990. 119(1): p. 25–30.

38. Briolat, V., et al., Contrasted Innate Responses to Two Viruses in Zebrafish: Insights into the Ancestral Repertoire of Vertebrate IFN-Stimulated Genes. The Journal of Immunology, 2014. 192(9): p. 4328–4341.

39. Ludwig, M., et al., Whole-body analysis of a viral infection: vascular endothelium is a primary target of infectious hematopoietic necrosis virus in zebrafish larvae. PLoS Pathog, 2011. 7(2): p. e1001269.

40. Bukreyev, A., et al., Nonsegmented negative-strand viruses as vaccine vectors. J Virol, 2006. 80(21): p. 10293–306.

41. Harmache, A., et al., Bioluminescence imaging of live infected salmonids reveals that the fin bases are the major portal of entry for Novirhabdovirus. J Virol, 2006. 80(7): p. 3655–9.

42. Souto, S., et al., In vivo multiscale analyses of spring viremia of carp virus (SVCV) infection: From model organism to target species. PLoS Pathog, 2024. 20(8): p. e1012328.

43. Casadevall, A., Climate change brings the specter of new infectious diseases. The Journal of Clinical Investigation, 2020. 130(2): p. 553–555.

44. Mora, C., et al., Over half of known human pathogenic diseases can be aggravated by climate change. Nature Climate Change, 2022. 12(9): p. 869–875.

45. Reid, G.K., et al., Climate change and aquaculture: considering biological response and resources. Aquaculture Environment Interactions, 2019. 11: p. 569–602.

46. Franke, A., A. Beemelmanns, and J.J. Miest, Are fish immunocompetent enough to face climate change? Biol Lett, 2024. 20(2): p. 20230346.

47. Alto, B.W., et al., Stochastic temperatures impede RNA virus adaptation. Evolution, 2013. 67(4): p. 969–79.

48. Sun, J., et al., Structure-based self-supervised learning enables ultrafast protein stability prediction upon mutation. Innovation (Camb), 2025. 6(1): p. 100750.

49. Aggad, D., et al., The two groups of zebrafish virus-induced interferons signal via distinct receptors with specific and shared chains. J Immunol, 2009. 183(6): p. 3924–31.

50. Walker, P.J., et al., ICTV Virus Taxonomy Profile: Rhabdoviridae 2026. J Gen Virol, 2026. 107(6).

51. Fijan, N., et al., Some properties of the Epithelioma papulosum cyprini EPC cell line from carp Cyprinus carpio. Annales de l’Institut Pasteur Virology, 1983. 134E: p. 207–20.

52. Fuerst, T.R., et al., Eukaryotic transient-expression system based on recombinant vaccinia virus that synthesizes bacteriophage T7 RNA polymerase. Proc Natl Acad Sci U S A, 1986. 83(21): p. 8122–6.

53. Bols, N.C., et al., Development of a cell line from primary cultures of rainbow trout, Oncorhynchus mykiss (Walbaum), gills. Journal of Fish Diseases, 1994. 17(6): p. 601–611.

54. Zamperin, G., et al., Sequencing of animal viruses: quality data assurance for NGS bioinformatics. Virology Journal, 2019. 16(1): p. 140.

55. Abramson, J., et al., Accurate structure prediction of biomolecular interactions with AlphaFold 3. Nature, 2024. 630(8016): p. 493–500.

56. Pettersen, E.F., et al., UCSF ChimeraX: Structure visualization for researchers, educators, and developers. Protein Sci, 2021. 30(1): p. 70–82.

57. Biacchesi, S., The reverse genetics applied to fish RNA viruses. Vet Res, 2011. 42(1): p. 12.

58. Alestrom, P., et al., Zebrafish: Housing and husbandry recommendations. Lab Anim, 2020. 54(3): p. 213–224.

59. Lister, J.A., et al., nacre encodes a zebrafish microphthalmia-related protein that regulates neural-crest-derived pigment cell fate. Development, 1999. 126(17): p. 3757–67.

60. Livak, K.J. and T.D. Schmittgen, Analysis of relative gene expression data using real-time quantitative PCR and the 2(-Delta Delta C(T)) Method. Methods, 2001. 25(4): p. 402–8.

61. Nguyen-Chi, M., et al., Transient infection of the zebrafish notochord with E. coli induces chronic inflammation. Dis Model Mech, 2014. 7(7): p. 871–82.

62. Guerra, J., et al., Lysyl-tRNA synthetase produces diadenosine tetraphosphate to curb STING-dependent inflammation. Sci Adv, 2020. 6(21): p. eaax3333.

63. Novoa, B., et al., Rag1 immunodeficiency-induced early aging and senescence in zebrafish are dependent on chronic inflammation and oxidative stress. Aging Cell, 2019. 18(5): p. e13020.

