## Supplemental tables for "Molecular characterization of thermo-adapted Novirhabdovirus variants and virulence *in vivo* in rainbow trout and zebrafish at different temperatures"

**Table S1. Distribution of mutations identified in the thermo-adapted IHNV_25_ among field isolates.**

| **IHNV_25_**  **Protein** | **Position** | **Field isolates^a^ (host species^b^/reported CPM in rainbow trout)** |
| --- | --- | --- |
| N | 108S | None |
| P | 14C | None |
|  | 169R | None |
| M | 104R | None |
| G | 67N | **Dfr858-07 (Om/NA)**, **IO3-14bw (Om/NA)**, **Dau2185-05 (Om/NA)**, **TV0203 (Om/NA)**, **RtJe00 (Om/NA)** |
|  | 98Y | **Dfr858-07 (Om/NA)**, **IO3-14bw (Om/NA)**, **Dau2185-05 (Om/NA)**, **TV0203 (Om/NA)**, **RtJe00 (Om/NA)**, **I/TN/13-2/Janv21 (Om/NA)**, I/BG/67-1/Jan22 (Om/94%), SA-22-05674 (Om/NA) |
|  | 178G | **I/TN/13-2/Janv21 (Om/NA)** |
| L | 702G | None |
|  | 872L | None |
|  | 1098C | None |
|  | 1523N | None |

^a^References of field isolates including the G gene sequence (GenBank accession number), the host species from which it has been isolated, and the cumulative percent of mortality reported from rainbow trout experimentally infected by bath immersion when available (NA: not available): Dfr858-07 (ACG50040), IO3-14bw (CUS19924), Dau2185-05 (ACG50035), TV0203 (BBA19181), RtJe00 (BAF64531), I/TN/13-2/Janv21 (WWQ80870), I/BG/67-1/Jan22 ([1]/ WWQ80862), SA-22-05674 ([2]/SUB16546936).

^b^Host species abbreviations: Om, *Oncorhynchus mykiss* (rainbow trout).

None: amino-acid substitutions absent from Genbank.

**Table S2. Distribution of mutations identified in the thermo-adapted IHNV_24_-Cherry among field isolates.**

| **IHNV_24_**  **Protein** | **Position** | **Field isolates^a^ (host species^b^/reported CPM in rainbow trout)** |
| --- | --- | --- |
| N | 391L | None^c^ |
| M | 104T | 3Lot1 (Om/NA), WRAC (Ot/84%), T00121 (Om/NA) |
| G | 55I | None |
|  | 57V | I/TN/200-2/Mar2023 (Om/NA), I/BG/67-1/Jan22 (Om/94%) |
|  | 78I | I01-1466 (Om/NA) |
| L | 408E | None |
|  | 441T | None |
|  | 1461T | None |
|  | 1831T | VP0614-8 (Om/NA), I/TN/91/Feb19 (Om/NA), I/TN/223/Mar18 (Om/NA), I/TN/62/Feb15 (Om/NA), I/TN/224/Mar18 (Om/NA), 00121-8 (El/NA) |

^a^References of field isolates including the M, G and L gene sequences (GenBank accession number), the host species from which it has been isolated, and the cumulative percent of mortality reported from rainbow trout experimentally infected by bath immersion when available (NA: not available): 3Lot1 (QGR26046), WRAC (NP_042678), T00121 (qgr26064), I/TN/200-2/Mar2023 (WWQ80872), I/BG/67-1/Jan22 ([1]/WWQ80862), VP0614-8 (QGR26073), I/TN/91/Feb19 (QPB44651), I/TN/223/Mar18 (QPB44651), I/TN/62/Feb15 (QGJ79631), I/TN/224/Mar18 (QPB44657), 00121-8 (QKO28324)

^b^Host species abbreviations: Om, *Oncorhynchus mykiss* (rainbow trout); Ot, *Oncorhynchus tschawytscha* (Chinook salmon); El, *Esox Lucius* (pike).

None: amino-acid substitutions absent from Genbank.

^C^only one sequence is available with a leucine in position 391 of the nucleoprotein (P19691). This sequence is the first sequence for IHNV N protein published by Gilmore and Leong [3] but its N terminal region is marked as “Compositionally biased region”.

**Table S3. Amino acid sequence identity and similarity between the proteins of IHNV CAN67 and IHNV 32-87.**

| **IHNV**  **Protein** | **% of identity** | **% of similarity** | **Number of aa substitutions / protein length** |
| --- | --- | --- | --- |
| N | 99.2 | 99.7 | 2/391 |
| P | 99.1 | 99.1 | 2/230 |
| M | 99.5 | 100 | 1/195 |
| G | 99.2 | 99.4 | 4/508 |
| NV | 98.2 | 99.1 | 2/111 |
| L | 99.8 | 99.8 | 4/1986 |

aa, amino acid

**Références**

1.	Marsella, A., et al., *Temperature impact on replication and virulence of European infectious hematopoietic necrosis viruses.* Aquaculture, 2025. **609**: p. 742786.

2.	Gourhannic, E., et al., *Epidemic Risks associated to high-temperature replicating infectious hematopoietic necrosis virus (IHNV) field isolates: from molecular characterization to host pathogen interactions.* Submitted, 2026.

3.	Gilmore, R.D., Jr. and J.A. Leong, *The nucleocapsid gene of infectious hematopoietic necrosis virus, a fish rhabdovirus.* Virology, 1988. **167**(2): p. 644-8.
