## Supplementary figures and images for "Molecular characterization of thermo-adapted Novirhabdovirus variants and virulence *in vivo* in rainbow trout and zebrafish at different temperatures"

### Supplemental Figure S1

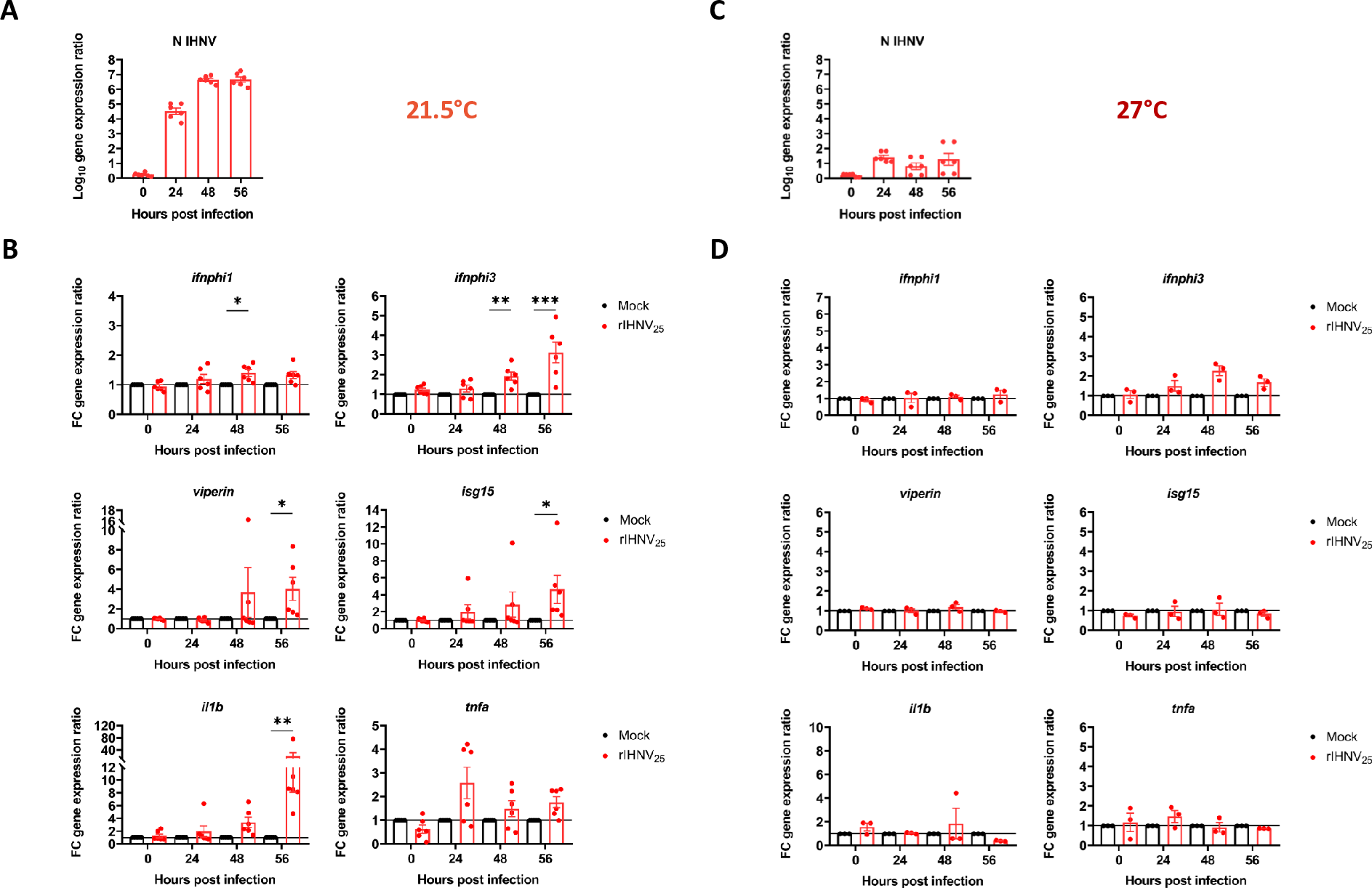

### Supplemental Figure S2

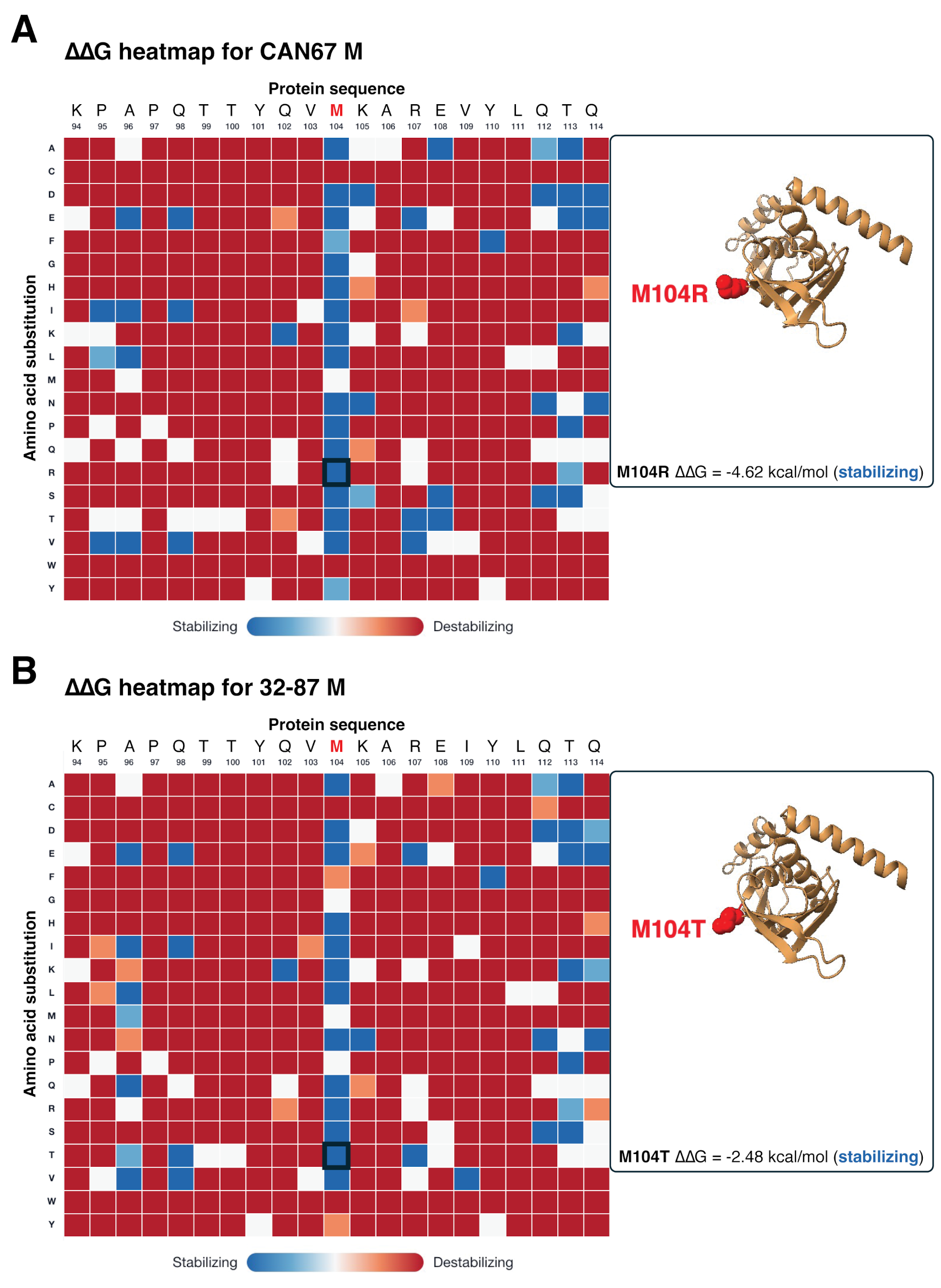
